# The haplotype-resolved chromosome pairs and transcriptome of a heterozygous diploid African cassava cultivar

**DOI:** 10.1101/2021.11.16.468774

**Authors:** Weihong Qi, Yi-Wen Lim, Andrea Patrignani, Pascal Schläpfer, Anna Bratus-Neuenschwander, Simon Grüter, Christelle Chanez, Nathalie Rodde, Elisa Prat, Sonia Vautrin, Margaux-Alison Fustier, Diogo Pratas, Ralph Schlapbach, Wilhelm Gruissem

## Abstract

**Background:** Cassava (*Manihot esculenta*) is an important clonally propagated food crop in tropical and sub-tropical regions worldwide. Genetic gain by molecular breeding is limited because cassava has a highly heterozygous, repetitive and difficult to assemble genome.

**Findings:** Here we demonstrate that Pacific Biosciences high-fidelity (HiFi) sequencing reads, in combination with the assembler hifiasm, produced genome assemblies at near complete haplotype resolution with higher continuity and accuracy compared to conventional long sequencing reads. We present two chromosome scale haploid genomes phased with Hi-C technology for the diploid African cassava variety TME204. Genome comparisons revealed extensive chromosome re-arrangements and abundant intra-genomic and inter-genomic divergent sequences despite high gene synteny, with most large structural variations being LTR-retrotransposon related. Allele-specific expression analysis of different tissues based on the haplotype-resolved transcriptome identified both stable and inconsistent alleles with imbalanced expression patterns, while most alleles expressed coordinately. Among tissue-specific differentially expressed transcripts, coordinately and biasedly regulated transcripts were functionally enriched for different biological processes. We use the reference-quality assemblies to build a cassava pan-genome and demonstrate its importance in representing the genetic diversity of cassava for downstream reference-guided omics analysis and breeding.

**Conclusions:** The haplotype-resolved genome allows the first systematic view of the heterozygous diploid genome organization in cassava. The completely phased and annotated chromosome pairs will be a valuable resource for cassava breeding and research. Our study may also provide insights into developing cost-effective and efficient strategies for resolving complex genomes with high resolution, accuracy and continuity.

## Background

High quality reference genomes are fundamental for genomic analyses, which have revolutionized the fields of biology and medicine. Most plant genomes are challenging to assemble with a high level of accuracy, continuity, and completeness because they vary in size, levels of ploidy and heterozygosity [1]. Particularly, many plant species, including cassava, can be clonally propagated, which can increase the effective number of alleles and heterozygosity [2–4]. Meanwhile, plant genomes are highly repetitive and contain abundant ancient and novel transposable elements [1, 5]. Intra-genomic heterozygosity and repeat elements are major sources of genome assembly errors [5, 6]. The cassava (*Manihot esculenta*) genome has a haploid genome size of 750 Mbp [7, 8], and is one of the most heterozygous [9] and repetitive [8] of currently sequenced plant genomes [10]. Despite continuous sequencing efforts using different technologies over the last decade, unresolved gaps and haplotypes persist in all chromosomes of currently available cassava genomes [7–9,11].

Cassava is an important staple crop that is clonally propagated in tropical and sub-tropical regions worldwide. The starchy storage roots are an important staple food for nearly a billion people and used for industrial purposes. In Africa, cassava is cultivated mainly by smallholder farmers because the crop produces appreciable yields under a wide array of environmental conditions. However, production is constrained by weeds, drought, pests, and most crucially, viral diseases. Therefore, breeding of more robust and productive cassava varieties is of high importance. Since conventional breeding of cassava is time-consuming, haplotype-resolved reference genomes with high continuity, accuracy and completeness will be a valuable resource for applications of genomic selection, genome editing and improving genetic gains in cassava breeding.

Continuous long reads (CLRs) produced by Pacific Biosciences (PacBio) Single Molecule, Real-Time (SMRT) sequencing technology and other long read sequencing technologies have been essential for generating reference quality genome assemblies cost effectively in the last decade [12]. The African cassava cultivars TME3 and 60444 have been sequenced and assembled using 70-fold PacBio CLRs (read N50 12 kbp), producing genome assemblies with contig N50 of 98 and 117 kbp, respectively [8]. Although both assemblies were much more continuous than all other reported cassava genomes [7, 9], they were still fragmented when measured by the continuity metric of a high-quality genome proposed by the Vertebrate Genome Project (VGP) consortium (contig N50 > 1 Mbp) [13]. Many haplotype alleles in TME3 and 60444 were reconstructed [8], but collapsed regions still persist throughout both assemblies because assembly of error-prone long sequencing reads (hereafter referred to as long reads) homogenized sequences from different haplotype alleles, paralogous loci and repeat elements [14]. The recently introduced PacBio high-fidelity (HiFi) sequencing technology is able to produce long (10-25 kbp) and highly accurate (>99.9%) sequencing reads (hereafter referred to as HiFi reads). For several human and animal genomes, equivalent or higher continuities have been achieved with HiFi reads [14–17]. Novel genome assemblers have been developed to leverage the full potential of HiFi reads [14, 18], where the combined performance of HiFi reads and HiFi-specific genome assemblers was benchmarked in assembling human and animal genomes. Their potential in assembling plant genomes is less well studied, but is gaining momentum [18, 19]. In comparison to the strawberry reference genome reconstructed from a combination of short Illumina sequencing reads and PacBio CLR [20], the HiFi assembly of Fragaria x ananass has contig N50 values that are 10 times higher. HiFi reads also enabled the assembly of the 35.6 Gbp California redwood genome [18]. The recently published haplotype-resolved potato genome [19] was generated using a combination of multiple sequencing strategies, including HiFi reads.

### Data description

In this study, we collected PacBio CLRs (ERR5487554 - ERR5487559), HiFi reads (ERR5485301), Illumina paired-end (PE) sequencing reads (hereafter referred to as Illumina PE reads) (ERR5484652), and Hi-C data (ERR5484651) for the African cassava cultivar TME204. It belongs to a group of cassava cultivars carrying the dominant monogenic CMD2 resistance locus, which provides resistance to Cassava Mosaic Diseases (CMD) caused by African Cassava Mosaic Viruses [21]. We benchmarked the performance of CLRs and HiFi reads in assembling this highly complex and heterozygous genome. Assembly continuity, accuracy, and haplotype resolution of different genome drafts produced by four CLR/HiFi assemblers [14,18,22] were evaluated using genome quality metrics proposed by the VGP consortium [23] with Illumina PE reads from the same sample. Our results demonstrate that HiFi reads are valuable in assembling a high-quality heterozygous and repetitive plant genome. The high base accuracy and long sequencing read length provide superior resolution and accuracy in resolving allele differences between haplotypes, paralogous genes and repeat elements. By combining HiFi reads with Hi-C data we produced a highly accurate, chromosome-scale, phased assembly for a diploid African cassava cultivar. The two haploid assemblies (PRJNA758616 and PRJNA758615) revealed extensive haplotype heterozygosity within a cassava diploid genome and provided for the first time a systematic view of the cassava diploid genome organization. To improve genome annotation, we further generated PacBio Iso-Seq reads (ERR5489420 - ERR5489422) from different tissues. The full-length transcript sequences identified not only novel transcripts and genes, but also revealed a highly complex transcriptome including expression of fusion transcripts and disrupted genes. The close to complete haplotype resolved, annotated genome also enabled pan-genome and allele-specific expression analysis, demonstrating the importance of a more complete representation of cassava genetic diversity for downstream reference-guided omics analysis and molecular breeding.

### Analyses

#### Cassava TME204 genome characteristics

Illumina PE reads (Table 1) were used to estimate the overall genome characteristics of TME204, revealing a highly heterozygous diploid genome different from the reference genome of the partially in-bred South-American cassava cultivar AM560 [7] and other well-studied genomes, such as the human reference genome. The peak for k-mers covering TME204 heterozygous sequence was as high as the peak corresponding to k-mers present in both haplotypes, while the k-mer coverage plots for the cassava and human reference genomes were dominated by their homozygous sequence peaks (Supplementary figure 1 a). Based on the number of variant-induced branches in the De Bruijn assembly graph, the level of heterozygosity in the TME204 genome was measured at 1%, which is a magnitude higher than the heterozygosity level in the cassava and human reference genomes (Supplementary figure 1 b). This value is a conservative estimate because it is based on genomic regions only with lower rates of nonstructural variations. Highly heterozygous regions introduce divergent paths with higher complexity, which cannot be resolved by conventional bubble calling algorithms used to calculate variant-induced branching rate [24]. Consequently, sequences with a high density of single nucleotide polymorphisms (SNPs), small insertions and deletions (indels ≤ 50 bp), and large structural variations (SVs, e.g. indels > 50 bp, duplications, inversions, and translocations) were not counted in the 1% of heterozygosity. The cassava genomes (TME204 and AM560) are more repetitive than the human reference genome (Supplementary figure 1 c), which makes them more difficult to assemble with high quality.

**Figure 1.**
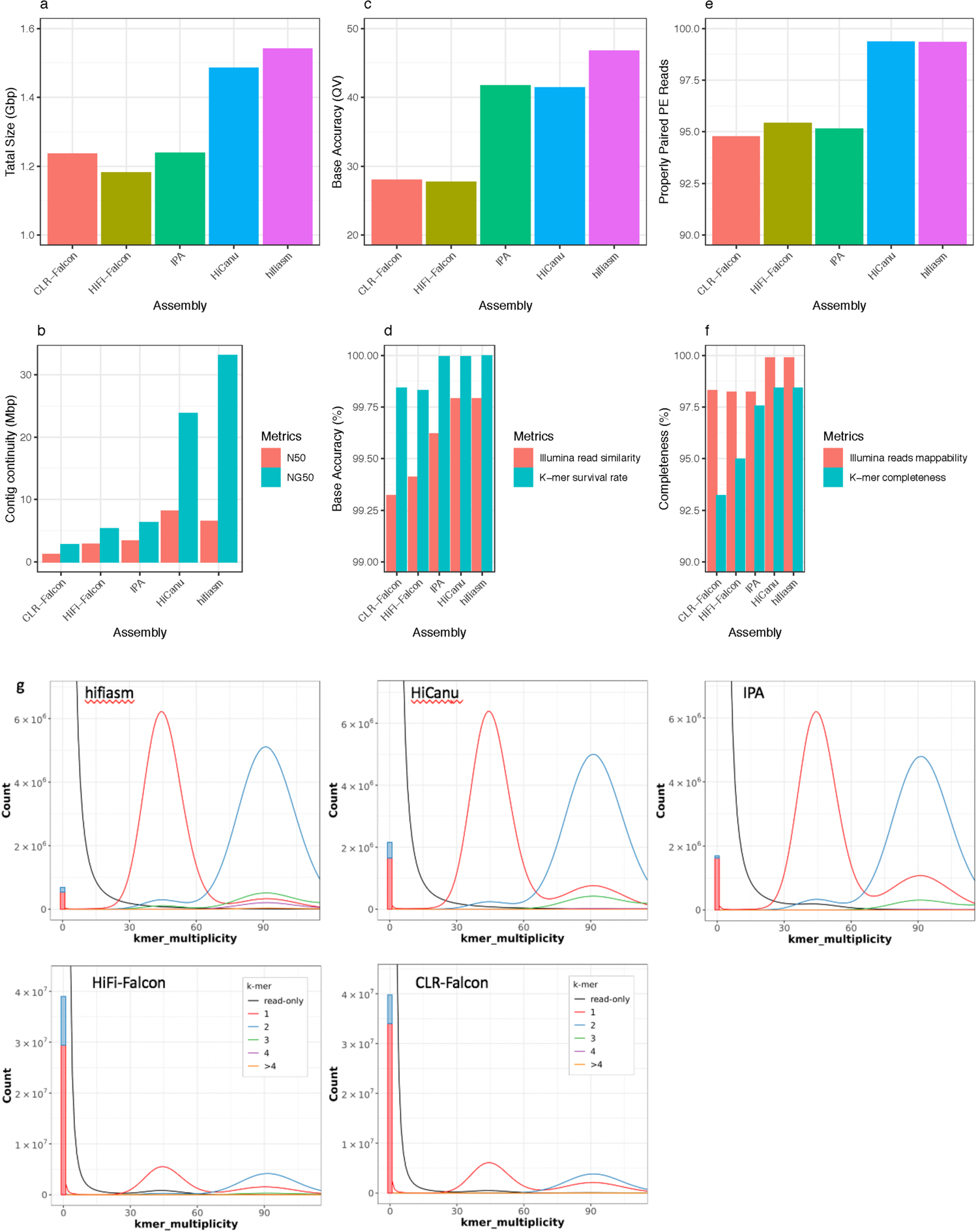
Benchmarking analysis of cassava TME204 assemblies from PacBio CLRs and HiFi reads. (a) Assembly size of the combined contig set of all resolved alleles. (b) Contig continuity measured as N50 and NG50. N50 is the length of the shortest contig in the set of largest contigs that make up 50% of the assembly size as shown in (a). NG50 is the length of the shortest contig in the set of largest contigs that make up 50% of the haploid genome size of 750 Mbp. (c) Phred scale of base accuracy of assembled sequences, calculated using the k-mer survival rate as shown in (d). (d) Base accuracy calculated using mapped Illumina reads and the fraction of k-mers found in the assembled sequences but missing in the Illumina reads, as shown in (g). (e) Structural accuracy of assembled sequences estimated using properly paired Illumina PE reads. (f) Assembly completeness estimated using the fraction of mapped Illumina reads and k-mer completeness (the fraction of reliable Illumina k-mers retained in the assembled sequences). (g) Merqury copy number spectrum plots of each assembly. K-mer coverage on the x-axis is computed from the Illumina reads. The y-axis is the abundance for k-mers with a given coverage, in the Illumina reads and the assembled sequences, respectively. K-mer was colored by the number of times they are found in the assembly. Homozygous k-mers found only once in the assembly (red hump at 80x) indicate collapsed haplotypes. Black humps found either at 40x (heterozygotes/1-copy k-mers) or 80x (homozygotes/2-copy k-mers) represent reliable Illumina k-mers missing in assembled sequences. The assembly specific k-mers absent from the Illumina reads are plotted as a bar at zero k-mer multiplicity.

**Table 1.**
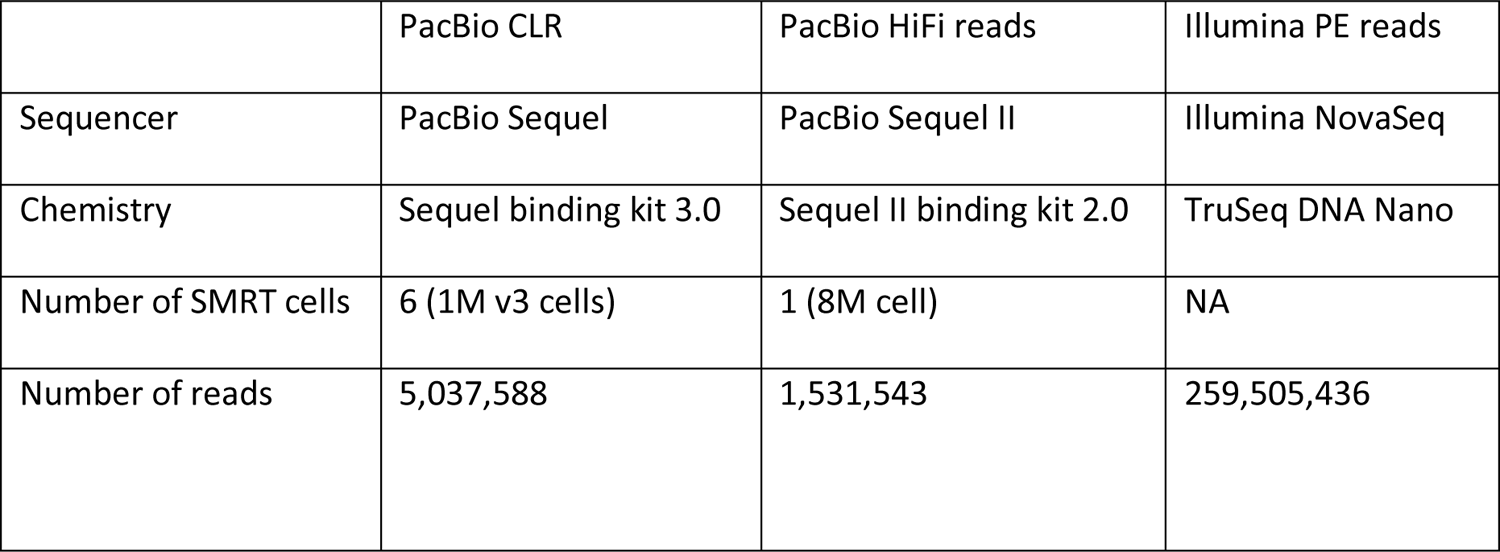
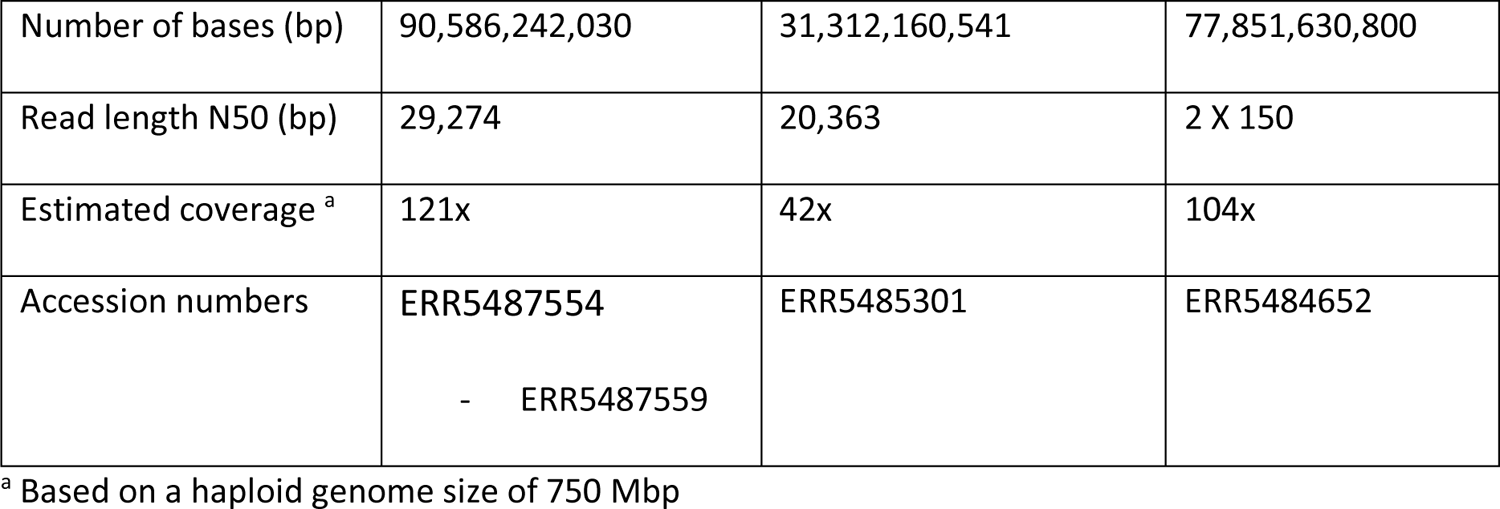
Cassava TME204 shotgun sequencing data collected.

|  | PacBio CLR | PacBio HiFi reads | Illumina PE reads |
| --- | --- | --- | --- |
| Sequencer | PacBio Sequel | PacBio Sequel II | Illumina NovaSeq |
| Chemistry | Sequel binding kit 3.0 | Sequel II binding kit 2.0 | TruSeq DNA Nano |
| Number of SMRT cells | 6 (1M v3 cells) | 1 (8M cell) | NA |
| Number of reads | 5,037,588 | 1,531,543 | 259,505,436 |
| Number of bases (bp) | 90,586,242,030 | 31,312,160,541 | 77,851,630,800 |
| Read length N50 (bp) | 29,274 | 20,363 | 2 X 150 |
| Estimated coverage <sup>a</sup> | 121x | 42x | 104x |
| Accession numbers | ERR5487554<br>- ERR5487559 | ERR5485301 | ERR5484652 |

<sup>a</sup> Based on a haploid genome size of 750 Mbp

#### Benchmarking cassava TME204 assemblies from PacBio CLR and HiFi reads

PacBio HiFi sequencing yielded 42x HiFi reads with length N50 of 20 kbp (Table 1). To assess the performance of different assemblers, the HiFi reads were assembled using four HiFi-specific software tools: Falcon, HiCanu, hifiasm and IPA (see Materials and Methods). For comparison of the HiFi reads with traditional long reads, we also assembled 121x PacBio CLRs (Table 1) from the same DNA sample using the Falcon assembler. In the case of a heterozygous, diploid genome such as cassava, diploid-aware assemblers (Falcon, hifiasm and IPA) produce a primary genome assembly, which is a set of primary contigs representing pseudo-haplotypes (the longest continuous stretches of assembled sequences), and an alternate assembly consisting of a set of unphased associated contigs (Falcon) or phased (IPA, hifiasm) haplotigs (i.e., continuous sequences of the same haplotype). For a genome with very high sequence divergence between haplotypes, haplotype allelic contigs can be incorrectly placed in the primary assembly, resulting in heterotype duplications that inflate the assembly size [23, 25]. The assembler hifiasm also identifies such contigs and places them into the alternate assembly. Phased haplotigs and purged contigs represent resolved alternative alleles (Supplementary table 1). In contrast to this strategy, HiCanu produces a single set of contigs representing all resolved alleles (Supplementary table 2).

**Table 2.** Assembly quality assessment of Cassava TME204 haplotigs.

| Quality Category | Quality Metric | Haplotype 1 | Haplotype 2 |
| --- | --- | --- | --- |
| General | Contig Size (Mbp) | 762 | 706 |
| Continuity | Contig N50 (Mbp) | 18 | 26 |
|  | Contig NG50 (Mbp) | 18 | 22 |
|  | Largest contig (Mbp) | 41 | 44 |
| Base accuracy | QV <sup>a</sup> | 45.23 | 48.94 |
|  | k-mer completeness (%) <sup>a</sup> | 79.6 | 79.1 |
| Structural<br>accuracy | k-mer false duplications (%) <sup>a</sup> | 3.0 | 2.4 |
|  | BUSCO duplicate (%) | 4.9 | 4.2 |
|  | Reference asmgene duplicate (%) | 1.6 | 0.8 |
|  | Reference gene transfer duplicate (%) <sup>b</sup> | 3.8 | 3.2 |
|  | Reliable blocks (%) | 97.2 | 97.5 |
|  | Congruent genetic markers (%) | 99.8 | 99.9 |
| Functional<br>completeness | BUSCO complete (%) | 99.0 | 98.8 |
|  | Reference asmgene complete (%) | 95.2 | 95.4 |
|  | Reference gene transfer rate (%) <sup>b</sup> | 96.3 | 96.5 |
|  | Transcript alignment rate (%) | 99.3 | 99.4 |

<sup>a</sup> QV and k-mer completeness for the combined assembly is 46.63 and 98.4%, respectively. The k-mer false duplication rate for the combined assembly is 3.5%.
<sup>b</sup> Calculated based on the number of lifted genes regardless of completeness

To achieve an unbiased comparison of the assembly continuity, accuracy, and completeness of all four assemblers, we first combined primary and alternate sequence assemblies into one contig set of mixed-haplotypes (Figure 1, Supplementary table 3). With the same amount of computing resources, HiFi read assembling was about two orders of magnitude faster and required ten times less data storage than CLR assembling. Each HiFi assembly was completed in only a few hours to a few days and used 20-800 GB of data storage when running on a single server with 64 CPUs and 500 GB RAM (random-access memory). CLR-Falcon assembly took a few weeks and used about 7 TB of disk space. The CLR-Falcon contig N50 of TME204 was already 10-fold longer compared to cassava TME3 and 60444 CLR-Falcon contigs [8], although the three genomes have similar levels of repetitiveness and heterozygosity (Supplementary figure 1). This could be due to longer sequencing read length, more accurate sequencing chemistry and base calling algorithms, higher coverage (Table 1), and the improved version of the Falcon assembler. HiFi reads improved contig continuity further, doubling contig N50 and NG50 values when assembled using Falcon (Figure 1 b). Assembled genome sizes varied based on the assembly software (Figure 1 a), thus N50 values cannot be used for comparisons between assemblers since they depend on the assembly size. Instead, we used NG50 [26] to compare assembly continuity among the HiFi assemblers, since it was normalized using the same haploid genome size of 750 Mbp instead of the varied total assembled sizes. According to NG50 values, the hifiasm contig set was the most continuous (NG50 33 Mbp), followed by the HiCanu contig set (23 Mbp) (Figure 1 b).

**Table 3.**
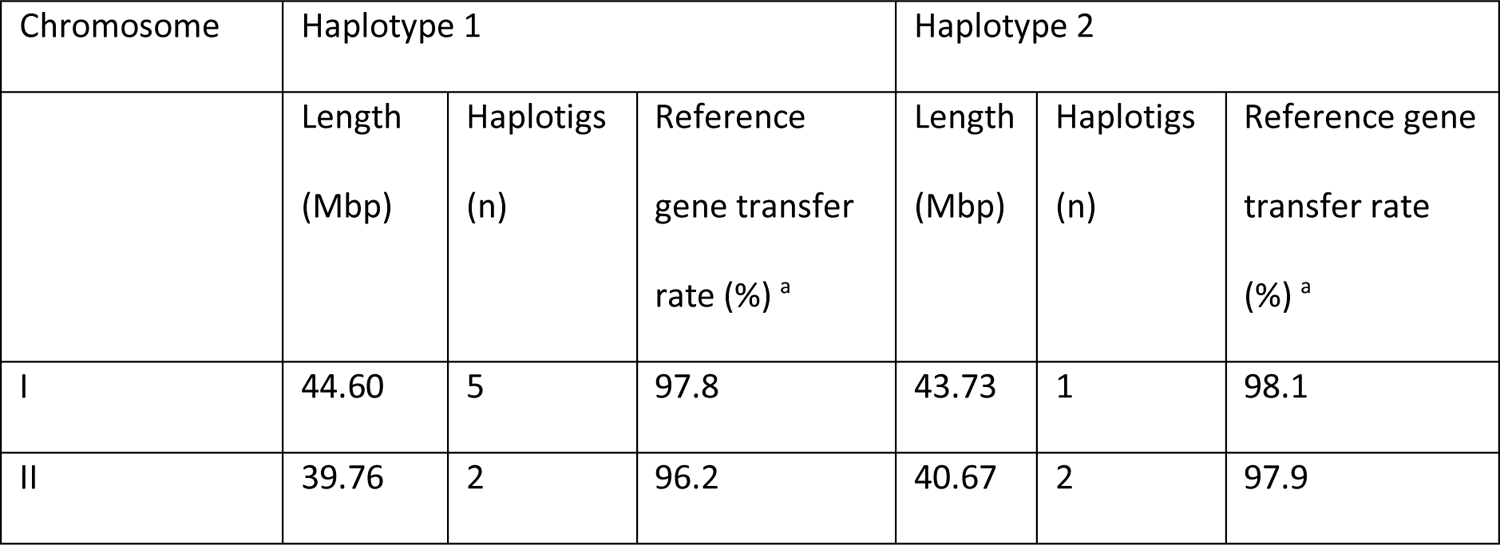
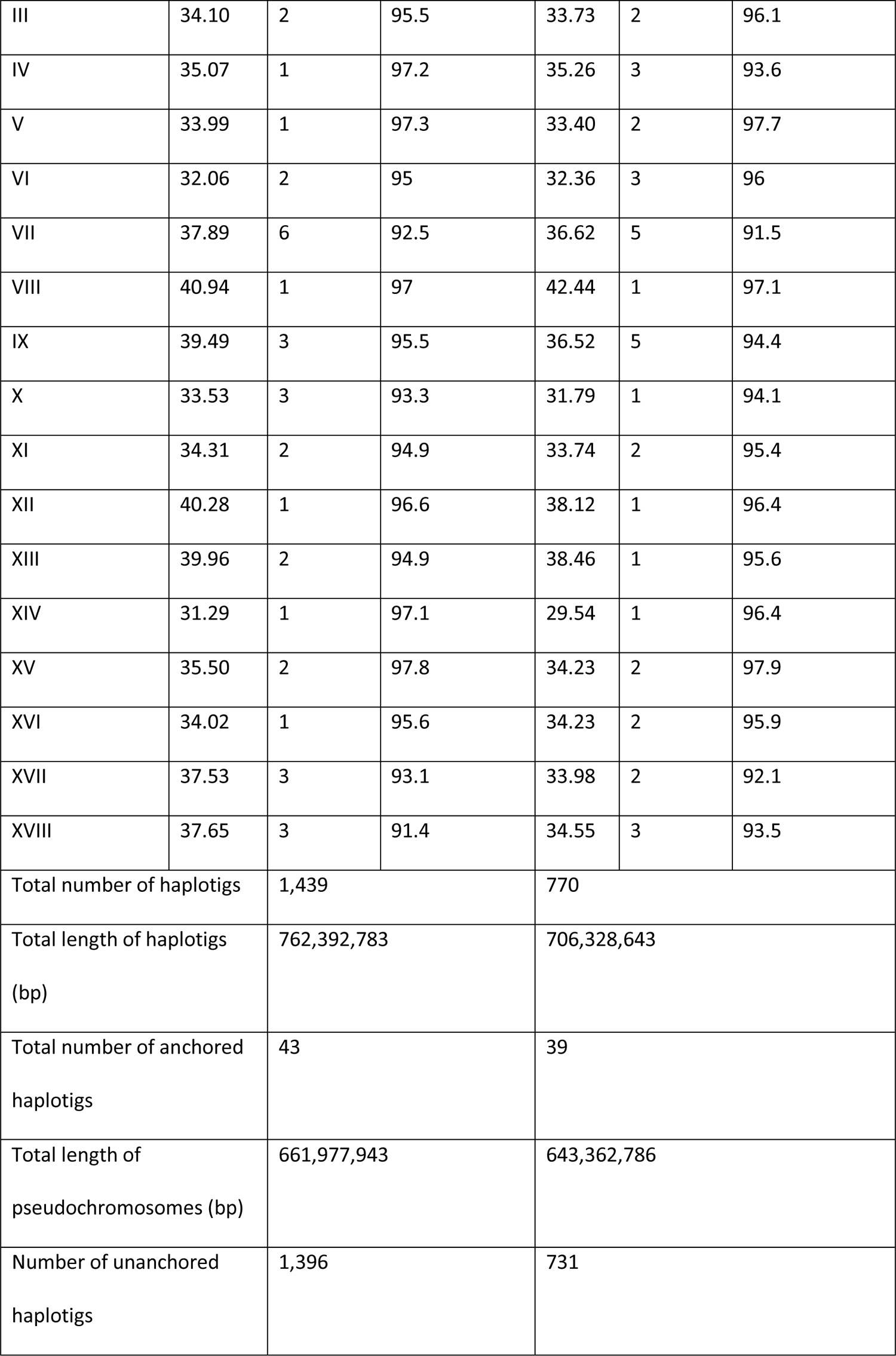
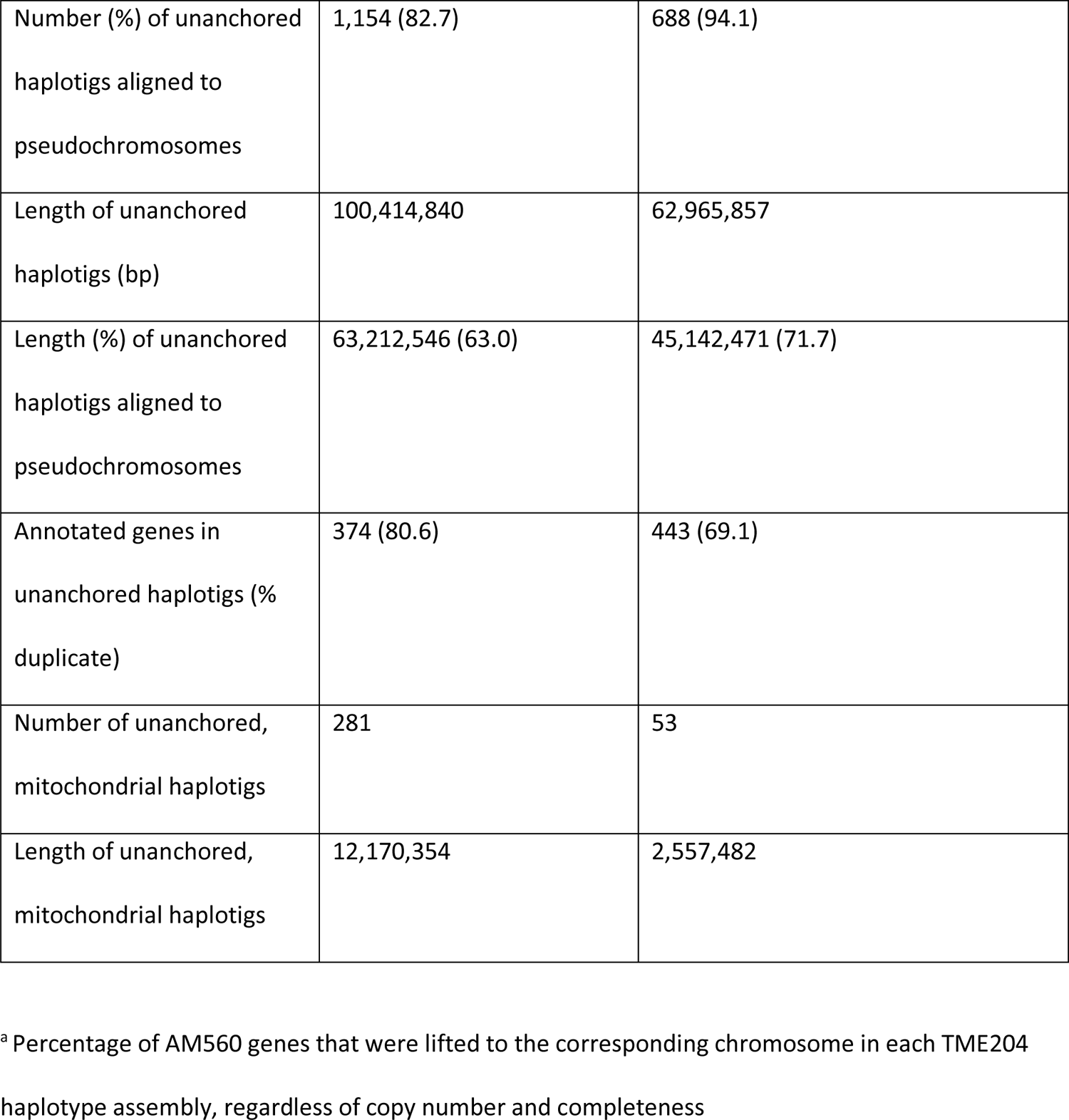
Phased chromosome pairs in TME204 diploid genome assembly.

| Chromosome | Haplotype 1 |  |  | Haplotype 2 |  |  |
| --- | --- | --- | --- | --- | --- | --- |
|  | Length<br>(Mbp) | Haplotigs<br>(n) | Reference<br>gene transfer<br>rate (%) <sup>a</sup> | Length<br>(Mbp) | Haplotigs<br>(n) | Reference gene<br>transfer rate<br>(%) <sup>a</sup> |
| I | 44.60 | 5 | 97.8 | 43.73 | 1 | 98.1 |
| II | 39.76 | 2 | 96.2 | 40.67 | 2 | 97.9 |
| III | 34.10 | 2 | 95.5 | 33.73 | 2 | 96.1 |
| IV | 35.07 | 1 | 97.2 | 35.26 | 3 | 93.6 |
| V | 33.99 | 1 | 97.3 | 33.40 | 2 | 97.7 |
| VI | 32.06 | 2 | 95 | 32.36 | 3 | 96 |
| VII | 37.89 | 6 | 92.5 | 36.62 | 5 | 91.5 |
| VIII | 40.94 | 1 | 97 | 42.44 | 1 | 97.1 |
| IX | 39.49 | 3 | 95.5 | 36.52 | 5 | 94.4 |
| X | 33.53 | 3 | 93.3 | 31.79 | 1 | 94.1 |
| XI | 34.31 | 2 | 94.9 | 33.74 | 2 | 95.4 |
| XII | 40.28 | 1 | 96.6 | 38.12 | 1 | 96.4 |
| XIII | 39.96 | 2 | 94.9 | 38.46 | 1 | 95.6 |
| XIV | 31.29 | 1 | 97.1 | 29.54 | 1 | 96.4 |
| XV | 35.50 | 2 | 97.8 | 34.23 | 2 | 97.9 |
| XVI | 34.02 | 1 | 95.6 | 34.23 | 2 | 95.9 |
| XVII | 37.53 | 3 | 93.1 | 33.98 | 2 | 92.1 |
| XVIII | 37.65 | 3 | 91.4 | 34.55 | 3 | 93.5 |
| Total number of haplotigs |  | 1,439 |  | 770 |  |  |
| Total length of haplotigs<br>(bp) |  | 762,392,783 |  | 706,328,643 |  |  |
| Total number of anchored<br>haplotigs |  | 43 |  | 39 |  |  |
| Total length of<br>pseudochromosomes (bp) |  | 661,977,943 |  | 643,362,786 |  |  |
| Number of unanchored<br>haplotigs |  | 1,396 |  | 731 |  |  |

|  |  |  |
| --- | --- | --- |
| Number (%) of unanchored haplotigs aligned to pseudochromosomes | 1,154 (82.7) | 688 (94.1) |
| Length of unanchored haplotigs (bp) | 100,414,840 | 62,965,857 |
| Length (%) of unanchored haplotigs aligned to pseudochromosomes | 63,212,546 (63.0) | 45,142,471 (71.7) |
| Annotated genes in unanchored haplotigs (% duplicate) | 374 (80.6) | 443 (69.1) |
| Number of unanchored, mitochondrial haplotigs | 281 | 53 |
| Length of unanchored, mitochondrial haplotigs | 12,170,354 | 2,557,482 |

<sup>a</sup> Percentage of AM560 genes that were lifted to the corresponding chromosome in each TME204 haplotype assembly, regardless of copy number and completeness

HiFi reads also improved accuracy and completeness of the assembled genome sequences. When measured using alignments of Illumina PE reads from the same sample (Table 1), both hifiasm and HiCanu achieved superior base accuracy (0.2% error rate) (Figure 1 d), structural accuracy (99.3% mapped reads were correctly paired) (Figure 1 e), and assembly completeness (99.9% mapped reads) (Figure 1 f). When measured using the Illumina k-mers analysis [27], the hifiasm assembly was most accurate (99.997%, quality value (QV) 46.74) and complete (98.40%) (Figures 1 c, d and g). The consensus accuracy measured using the aligned Illumina PE reads was lower than the measurement using k-mer analysis, while completeness measured using mapped Illumina PE reads was higher. The sequence differences between the mapped Illumina PE reads and the assembled consensus sequences could originate from sequencing errors in the Illumina reads and/or mis-alignments introduced by the alignment software tool. The read alignment process is also more tolerant to sequencing errors than the k-mer analysis, where erroneous k-mers were identified and excluded. Consequently, hereafter we only used k-mer analysis [27] to measure the accuracy and completeness of the genome sequence assemblies.

Because a correct assembly of PacBio CLR reads requires signal level polishing to achieve satisfactory consensus accuracy [28], we further phased and polished the CLR-Falcon assembly using Falcon-Unzip [22]. The final CLR-Falcon-Unzip assembly had a 39.75 and 34.22 QV score for the primary contigs and phased haplotigs, respectively. The QV score for the combined contig set was 38.86, with k-mer completeness of 97.64%. The numbers from all measurements were still worse than those achieved by HiCanu and hifiasm with HiFi reads, while using much less computing time and resources.

Similar to previous observations in TME3 and 60444 genome assemblies [8], all assemblers produced a total genome assembly of 1.2 Gbp or larger for TME204 (Figure 1 a, Supplementary table 2). All primary assemblies resolved with default parameter settings were also larger than the cassava haploid genome size estimated at 745 – 768 Mbp based on flow cytometry [8]. The BUSCO duplication rates varied but remained high (Supplementary table 1), underlining the difficulty of assembling the highly heterozygous and repetitive cassava genome. The total size of TME3 and 60444 assemblies could be decreased to around 750 Mbp with purge_haplotigs [25], a tool that identifies and removes allelic variants in the primary assembly, confirming that the large assembly size was mainly caused by heterotype duplications [8]. For the HiCanu and hifiasm TME204 assemblies, the total sizes were both about twice the haploid genome size, with more than 80% of the single-copy orthologous genes in the BUSCO plant database being duplicated (Supplementary table 2), indicating that they captured alleles from both haplotypes. Falcon (HiFi and CLRs) and IPA produced smaller assemblies, with lower numbers of duplicated BUSCO genes, further indicating they collapsed more haplotype alleles. Merqury k-mer analysis [27] confirmed that the assemblers varied in their performance of resolving haplotypes in the TME204 genome. The analysis identified hifiasm as the assembler producing the most haplotype-resolved assembly (Figure 1 g).

### Phased, haplotype-resolved contigs of cassava TME204

Subsequently we used a newer release of the hifiasm assembler (v0.15.2) to assemble the haplotype-resolved TME204 contigs, which were also phased at the same time using Hi-C technology (Table 2). The resulting two sets of haplotigs (phased haplotype-resolved contigs) represent haplotype 1 (762 Mbp) and haplotype 2 (706 Mbp) of the diploid cassava genome, hereafter referred to as H1 and H2, respectively. The assembly has a QV score of 45.23 for H1 haplotigs and 48.94 for H2 haplotigs, and 46.63 for the combined set of diploid contigs. For each haplotype and the combined diploid assembly, the k-mer completeness is 79.6%, 79.1%, and 98.4%, respectively, indicating that about 19% of the k-mers were haplotype-specific. Most importantly, k-mer analysis revealed that most haplotype-specific k-mers are present only once in the assembled sequences, while homozygous k-mers shared by two haploid genomes are present twice (Figure 2). This would be expected for a completely haplotype-resolved genome assembly in which even homozygous segments of the genome are included in both haplotypes. Only 3% of k-mers were from artificial duplications (Figure 2), which was similar to the false duplication rates measured at approximately 1% (reference asmgene score) to 4% (BUSCO duplication score). Functional completeness measured using plant BUSCO orthologs and TME204 Iso-Seq transcripts was 98% and above. The reference asmgene completeness score was slightly lower (95%). The slightly lower asmgene scores could be due to the high level of sequence differences between AM560 and TME204 (see later comparative analysis), therefore fewer AM560 reference genes could be aligned to TME204, resulting in lower asmgene completeness and duplication scores.

**Figure 2.**
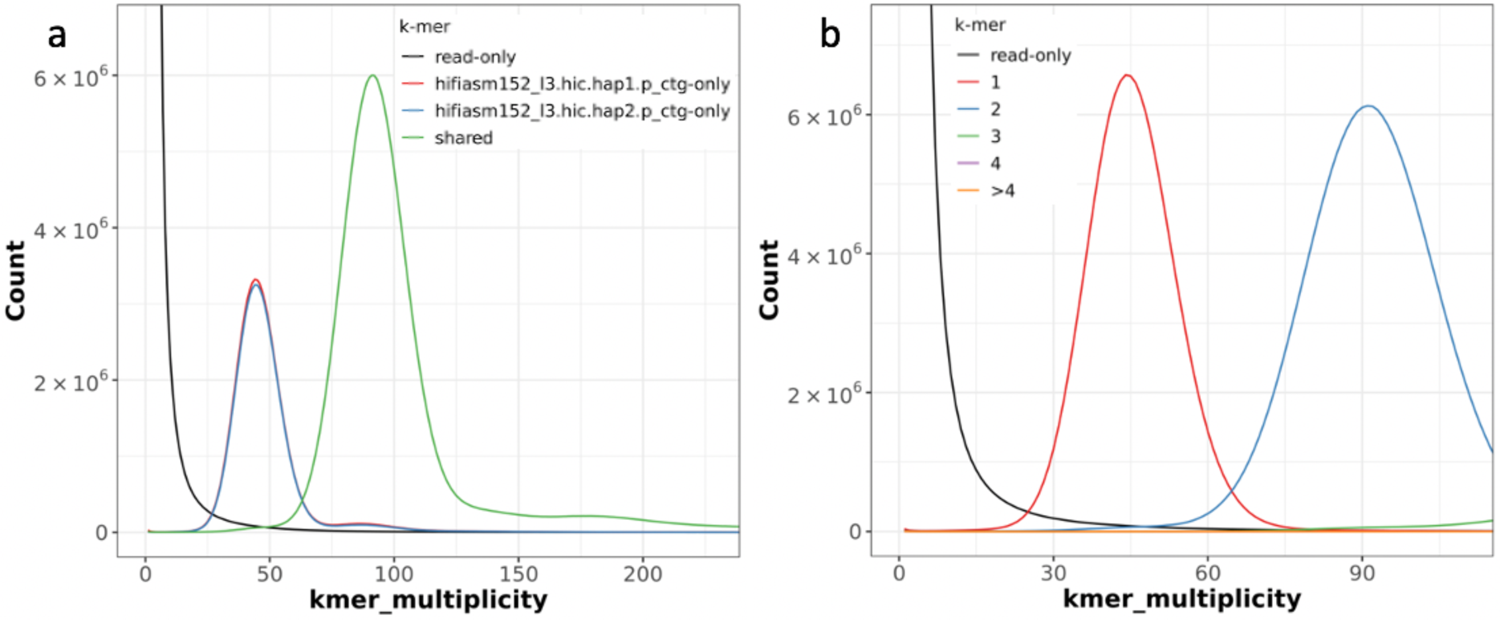
Merqury assembly and copy number spectrum plots of TME204 haplotigs. (a) For the Merqury assembly plot, k-mers are colored by their uniqueness in the Illumina PE reads (black), HiFi primary (red) and alternate (blue) assemblies. Shared k-mers are shown in green. At the heterozygotes peak (45 x), the second haplotype has only slightly fewer sequences (blue) compared to the first haplotype (red), indicating the reconstruction of heterozygous variants was almost complete. Red hump and blue shoulder around 90x are haplotype specific k-mers that are actually from homozygotes sequences, green shoulder around 45 x is due to shared k-mers belonging to heterozygotes. These shoulders are all very small, suggesting a very low level of collapsed homozygous regions and artificial duplications. (b) In the copy number spectrum plot, the majority of heterozygous k-mers appear once (red peak at 45x) and the majority of homozygous k-mers twice (blue peak at 90x) in the copy number spectrum plot, confirming that the assembly is close to complete haplotype-resolved and even the homozygous part of the genome is included in both haplotypes. High k-mer completeness is supported by the lack of black humps at 45x or 90x. Low artificial duplication is revealed by the barely detectable humps (green, purple, orange) of duplicated k-mers. The bars at zero k-mer multiplicity are low in both plots, suggesting most k-mers in the assemblies are also present in Illumina reads and therefore the assembled sequences are of high consensus accuracy.

To assess the structural accuracy of the assembled TME204 haplotigs, we mapped the longer PacBio CLR sequences from the same DNA sample (Table 1) to the haplotigs and analyzed the CLR read coverage along each haplotig (Supplementary figure 2). We defined reliably assembled sequences as those with at least 10x CLR read coverage. More than 97% of the assembled bases could be classified as correctly assembled with this quality metric.

To further validate the base, structural, and phasing accuracy of TME204 haplotigs, we generated complete sequences (96 to 128 kbp) of bacterial artificial chromosomes (BACs) containing TME204 genome fragments and aligned them to both sets of haplotigs (Figure 3, Supplementary figure 2). When a region is properly assembled and phased, we expect one continuous BAC-to-haplotig alignment for the corresponding BAC (resolved BAC). Three of the four sequenced BACs were resolved in H1 and one was resolved in H2, either perfectly or with only one indel difference (Supplementary table 4), confirming the close to Q50 consensus accuracy (i.e. one error per 100 kbp consensus sequences). The striking differences of BAC-to-haplotig alignments between the two haplotypes highlight the high level of haplotype differences in these regions.

**Figure 3.**
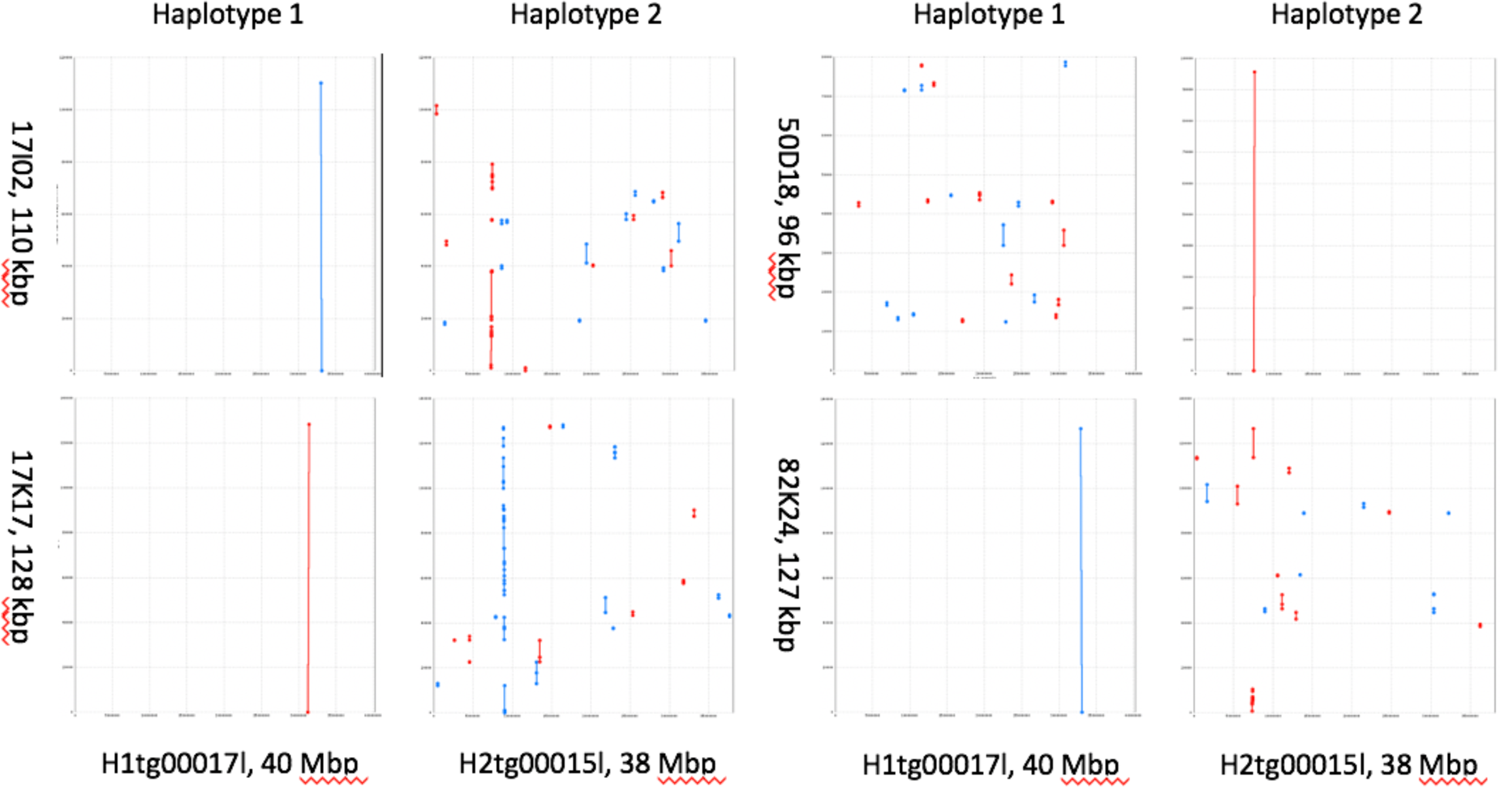
Phasing accuracy of TME204 haplotigs validated using BAC-to-haplotig alignments. Each dot plot shows the alignment of one BAC (y-axis) with one haplotig (x-axis). Forward alignments are plotted as red lines/dots, reverse alignments in blue. A line represents an undisturbed segment of alignment. When a region is correctly assembled and phased, the corresponding BAC sequence will align continuously (a resolved BAC). Three of the four BACs were resolved in the TME204 H1 assembly, the fourth was resolved in the TME204 H2 assembly. For each BAC, the striking differences of BAC-to-haplotig alignments reveal the high level of haplotype differences in these regions.

**Table 4.** Inter- and intra-genomic diversity of cassava revealed by comparative analysis of assembled contig sequences.

|  | Inter-genomic | Intra-genomic |
| --- | --- | --- |
| Comparison | TME204 H1 vs. AM560 | TME204 H1 vs H2 |
| Too divergent to be aligned (Mbp) | 220 (29%) | 181 (24%) |
| Uniquely aligned (Mbp) | 387 | 420 |
| Sequence similarity (%) in uniquely aligned regions | 98.75 | 98.85 |
| Number of SNPs | 2,720,699 | 2,720,467 |
| Number of single nucleotide indels | 1,874,181 | 1,867,232 |
| Number of Assemblytics small indels (20 – 50 bp)<br>(% expansion/contraction of repeats) | 13,605 (3%) | 13,332 (3%) |
| Number of Assemblytics large indels (50 bp – 10<br>kbp) (% expansion/contraction of repeats) | 13,387 (67%) | 13,213 (67%) |

The haplotigs were also compared with the cassava high-density genetic map [29]. Among the 22,403 available genetic makers, about 14,000 could be uniquely aligned to each set of the TME204 haplotigs with full length coverage and 100% sequence identity. More than 99.8% of these unique genetic markers showed high congruence between the genetic map and assembled haplotigs (Supplementary figure 3). Only less than 0.2% of the genetic markers were found among markers with different chromosome origins. Plots of genetic versus physical distance identified three pairs of chromosome-scale haplotigs (chromosomes VIII, XII and XIV) and six other chromosome-scale haplotigs in either H1 or H2 (Table 3). In plots of genetic versus physical distance for these chromosome-scale haplotigs (Supplementary figure 3), we often observed steep slopes at the haplotig ends and flat regions in their centers, which is consistent with increased recombination in chromosome arms and reduced recombination in pericentric regions of the chromosomes. Collectively, the data suggest that all of the 18 cassava chromosome pairs are highly continuous at the haplotig level and composed of only a few haplotigs per chromosome.

### Pseudochromosome pairs of cassava TME204

To further scaffold haplotigs into pseudochromosomes, we first used Hi-C scaffolding, but this did not further scaffold any haplotigs in H1 (Supplementary File 1). In H2, Hi-C data produced seven chromosomal scaffolds that were perfectly congruent with the genetic map, but also mis-joined haplotigs from different chromosomes (Supplementary table 5, Materials and Methods). Together, the high congruence between haplotigs and the genetic map allowed us to reconstruct all 18 pairs of pseudochromosomes with high confidence (Figure 4, Supplementary File 2). TME204 H1 and H2 pseudochromosomes are composed of 43 and 39 haplotigs, respectively. In total 12 pseudochromosomes are chromosome scale (Table 3). Haplotig orientations could be determined (Supplementary file 3) except for two small haplotigs in H2, representing the first 0.4 Mbp of chromosome VII and the last 1.3 Mbp of chromosome XI (Supplementary file 4). Together, 86.8% and 91.1% of the haplotig sequences could be assigned to chromosomes for H1 and H2, respectively (Table 2, Supplementary Files 3 and 4).

**Figure 4.**
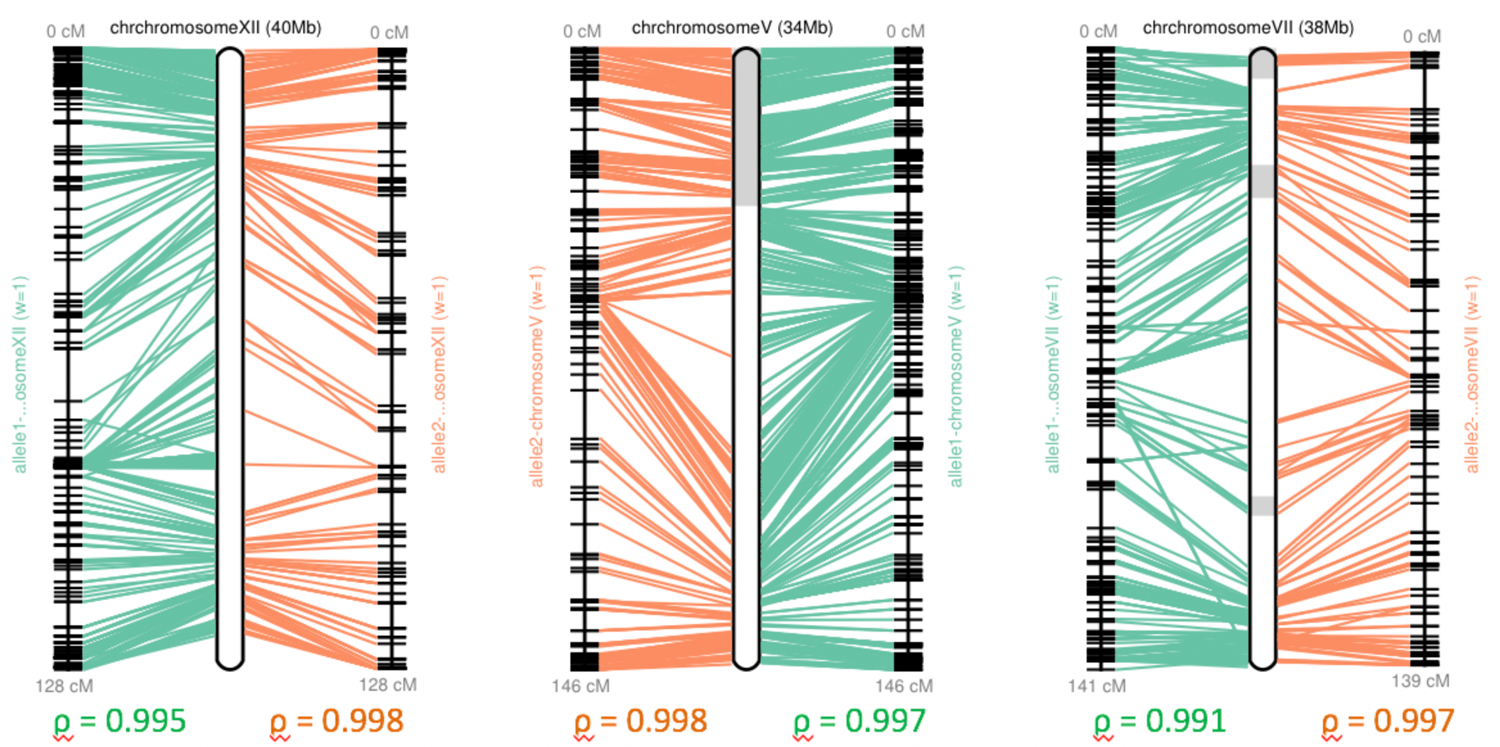
Reconstruction of pseudochromosomes in the cassava TME204 H1 assembly using the high-density genetic map. For each pseudochromosome, the panel shows the physical positions on the reconstructed pseudochromosome and the map positions connecting by lines. Adjacent contigs within the reconstructed pseudochromosome are shown as boxes with alternating shades. The ρ-value under each map measures the Pearson correlation coefficient, with values in the range of −1 to 1, and values closer to −1 and 1 indicate near-perfect collinearity. Chromosome XII is composed of a single chromosomal haplotig, the same as for chromosomes IV, VIII, XIV, and XVI. Chromosome V is composed of two contigs, the same as for chromosomes II, III, VI, IX, X, XI, XIII, XV, XVII, XVIII. Chromosome VII is composed of six haplotigs, which is the most fragmented chromosome in the TME204 H1 assembly, followed by chromosome I that has four haplotigs.

In both TME204 H1 and H2 assemblies, we found haplotigs that could not be scaffolded using either the genetic map (Table 3) or Hi-C technology (Supplementary Files 1 and 2). A majority of these unanchored haplotigs can be partially aligned to the pseudochromosomes with an average sequence similarity of 98% (Table 3). A few hundred AM560 genes can be transferred onto these haplotigs as well, although most (70%) were duplicated copies of genes that already transferred onto pseudochromosomes. It is clear that these haplotigs are of cassava origin and not from foreign contamination. When the assembled sequences were screened against the NCBI (National Center for Biotechnology Information) mitochondrial database, unanchored haplotigs representing the highly fragmented mitochondrial genome were identified in both haplotype assemblies (Table 3, Supplementary figure 4 a). When compared to the other none mitochondrial unanchored haplotigs, mitochondrial haplotigs have a smaller size variation (25-76 kbp) and lower depth of coverage on average (Supplementary figure 4 b). Regions similar to nuclear mitochondrial pseudogene regions (numt’s) were also ubiquitous and found in both pseudochromosomes (Supplementary figure 4 c) and unanchored haplotigs (Supplementary table 6). Some of the none-mitochondrial unanchored haplotigs can be regions still missing from the current set of pseudochromosome pairs where the gene content completeness ranges from 91 to 98% (Table 3). They can also be results of assembly artifacts (i.e. collapsed repeats) or represent novel haplotypes from *de novo* mutations.

### Repeat and gene landscape of cassava TME204 genome

*De novo* repeat modeling using all resolved allelic sequences (i.e. primary plus alternate contigs) identified 1,431 repeat families that make up 20% of TME204 genome, with 1,016 families representing novel unclassified repeats. The distribution of family sizes and sequence lengths among the novel repeat families is similar to those in LTR families, which masked up to 40% of TME204 genome. In total, 69% of each TME204 haploid genome is masked as repeats (Supplementary figure 5), which is slightly higher than the level reported for TME3 and 60444 genomes (65%) [8].

**Figure 5.**
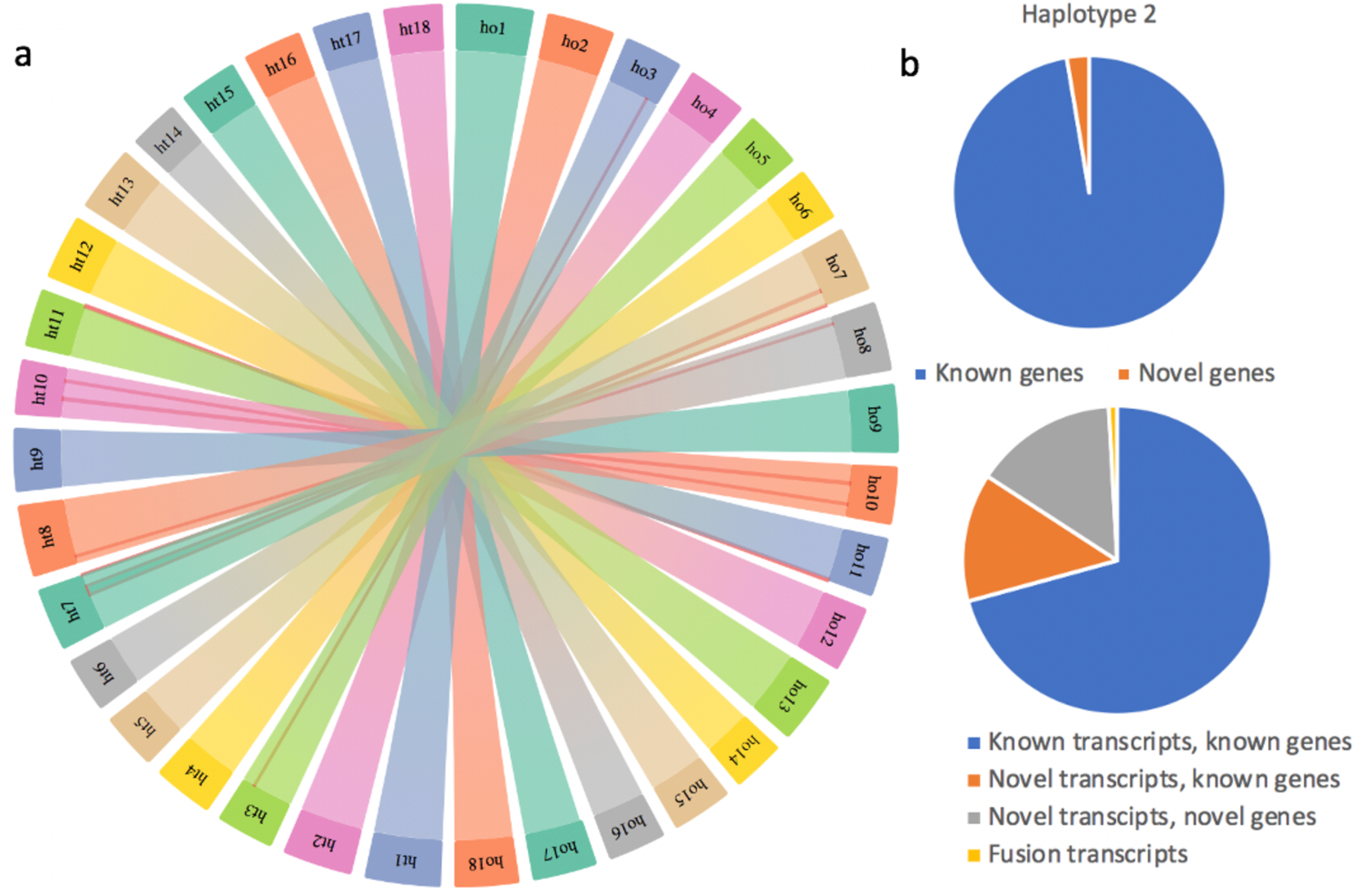
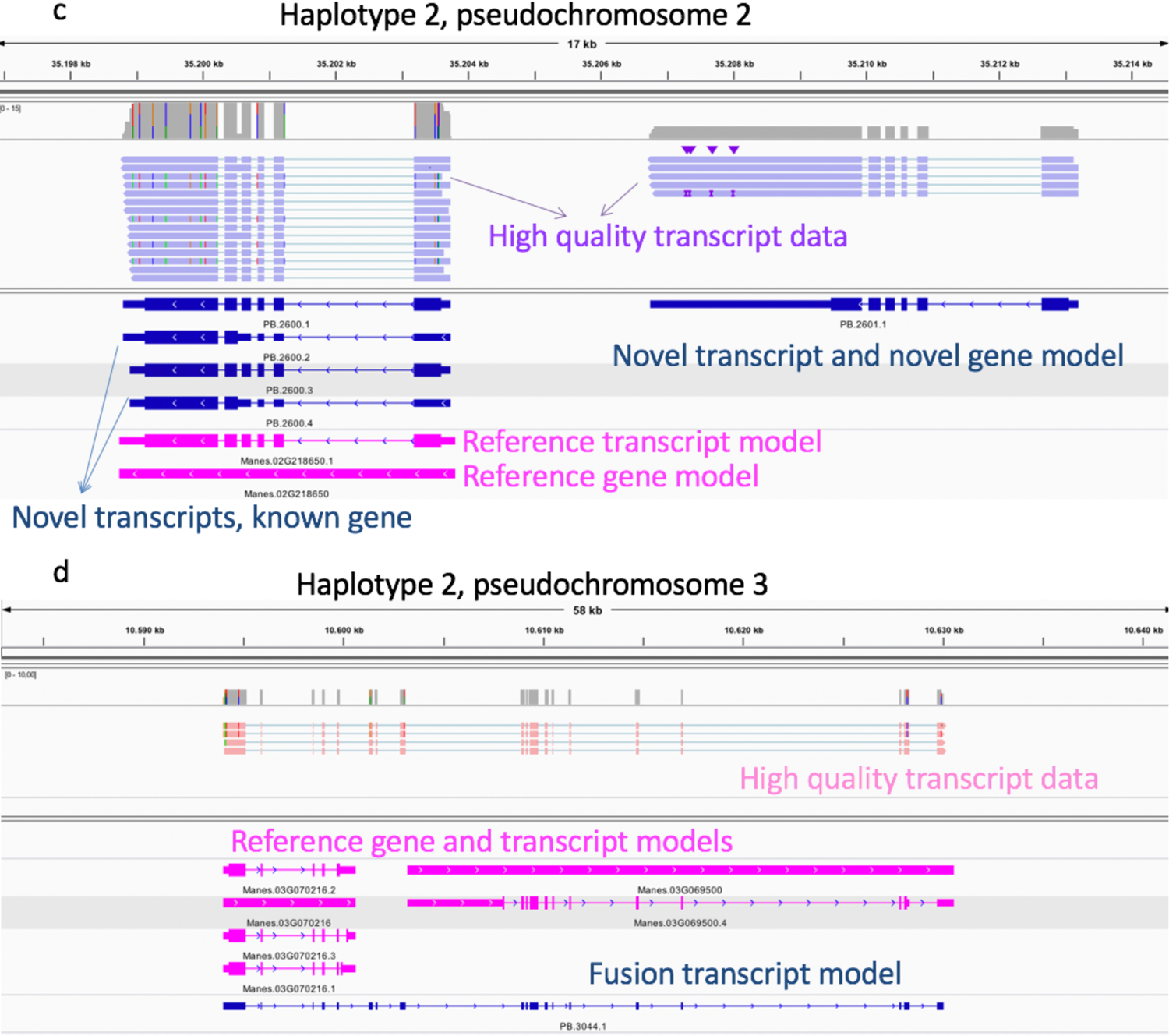
Validation and improvement of TME204 gene annotation using PacBio Iso-Seq transcripts. (a) Gene synteny between the two TME204 pseudochromosome pairs revealed by orthologous pairs of lifted genes. “ho” and “ht” encode “haplotype 1” and “haplotype 2”, respectively. Color lines highlight the inverted regions. (b) Lifted genes/transcripts validated and novel genes/transcripts identified using PacBio Iso-Seq transcripts. (c) Example of a novel transcript (PB.2601.1) from a novel gene model, which is found next to a validated reference gene model (Manes.02G218650) with two validated transcripts (PB.2600.2 and PB.2600.4) and two novel transcripts (PB.2600.2 and PB.2600.4). (d) Example of a fusion transcript spanning two reference gene models (Manes.3G070216 and Manes.03G069500) on pseudochromosome III in the TME204 H2 assembly.

In the last 10 years, continuous efforts have been made to improve the assembly and annotation of the cassava reference genome AM560 [7,11,29,30]. The set of AM560 reference gene models (https://phytozome-next.jgi.doe.gov/info/Mesculenta_v8_1) is widely used in the research field. Since the AM560 genome is from an inbred South-American cassava cultivar, our high-quality haplotype-resolved genome of the heterozygous African TME204 cultivar supplements AM560 as a reference for other heterozygous cassava genomes. We therefore annotated the TME204 genome by transferring well established cassava reference gene models, including CDSs (coding sequences), transcripts/mRNAs, and genes, to TME204 H1 and H2 assemblies. Over 97% of the 32,805 AM560 genes could be lifted to each TME204 haplotype assembly with a duplication rate of 3 to 4%, which is similar to BUSCO complete and duplicate scores (Table 2). Among the transferred genes, 9% (2,821 and 2,790 genes in H1 and H2, respectively) appeared as disrupted protein coding genes because all the associated transcripts were disrupted or incomplete after being lifted. While some of the disrupted protein coding genes became incomplete in both TME204 haplotypes, others were found incomplete in one haplotype only (Supplementary figure 6 a). The functional implication of these haplotype-specific incomplete transcripts was investigated using Gene Ontology (GO) enrichment analysis. Although some of the enriched GO terms in the category of biological process (BP), such as phosphorylation and protein phosphorylation, are common for both sets of incomplete transcripts, others are haplotype-specific. Among H2-specific incomplete transcripts, GO terms related to the reproduction process, such as pollination, recognition of pollen etc. were enriched. For H1-specific incomplete transcripts, GO terms related to carbohydrate metabolism process and different catabolic processes were enriched (Supplementary figure 6 b).

**Figure 6.**
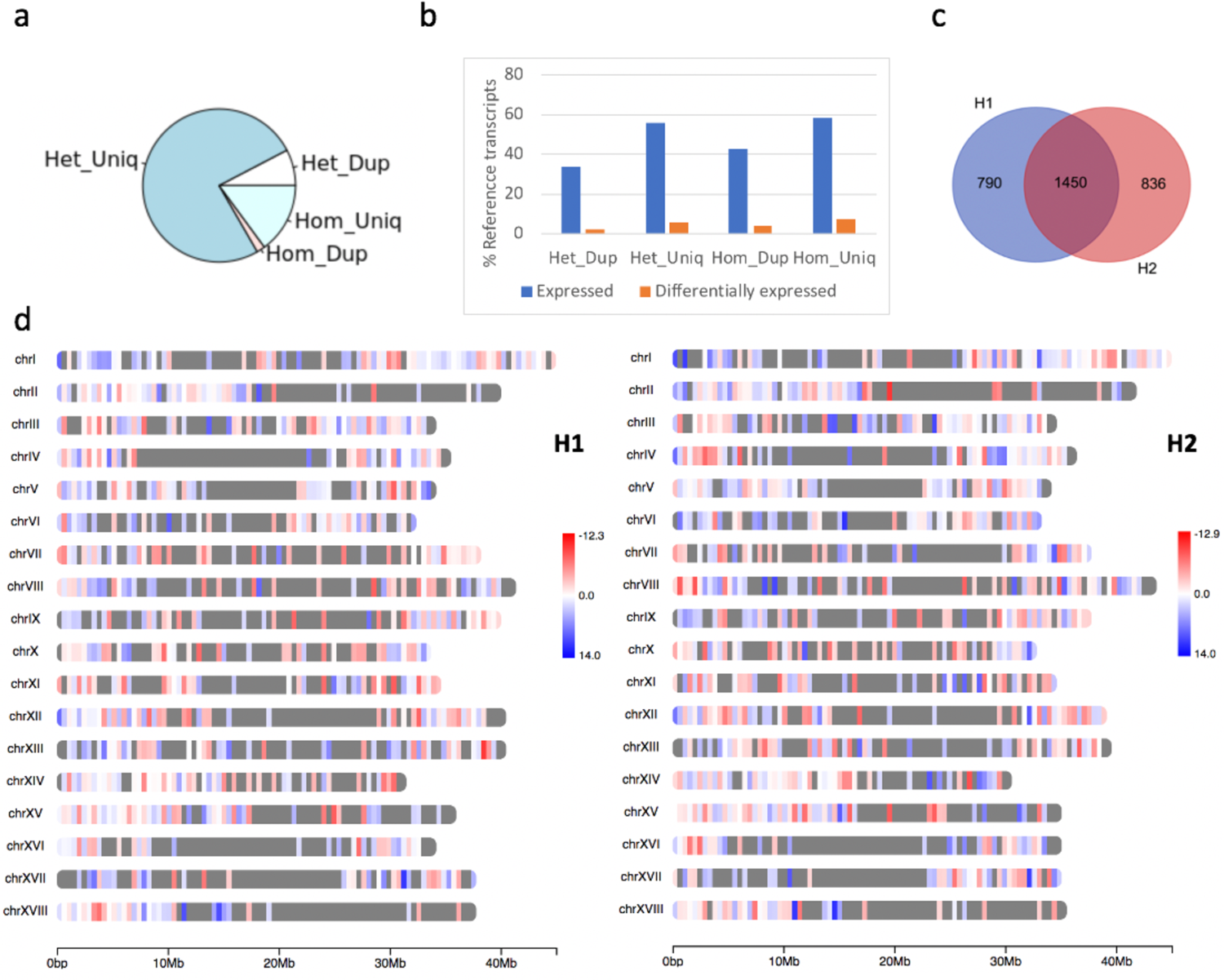
Haplotype resolved transcriptome analysis of TME204 transcripts differentially expressed between leaf and stem tissues. (a) The cassava TME204 haplotype resolved transcriptome is mainly composed of transcripts with different sequences between haplotypes. (b) Multi-copy transcripts are less highly expressed and differentially regulated (adjusted p-value < 0.001 and fold changes > 4) between TME204 leaf and stem tissues. (c) Overlapping of H1 and H2 genes where associated transcripts were differentially expressed between TME204 leaf and stem tissues. (d) Distribution of tissue specific differentially expressed transcripts on pseudochromosomes of TME204 H1 and H2. Color scales represent log2 fold changes. The transcriptome comparison between TME204 leaf and stem tissues identified gene loci with associated transcripts that were differentially regulated in one haplotype only.

Excluding these disrupted genes, 34,881 and 34,980 protein coding genes with 53,370 and 53,295 transcripts were annotated in the TME204 H1 and H2 assembly, respectively. Comparison of 26,602 chromosomal orthologous gene pairs revealed high gene synteny (99.08%) between TME204 pseudochromosome pairs. Only seven inverted regions involving 294 genes were detected on pseudochromosomes III, VII, VIII, X, and XI (Figure 5a). Gene synteny is also highly conserved between pseudochromosomes of AM560 and TME204 H2, where 98.69% of 27,908 orthologous gene pairs are kept in the same order, with nine inverted regions (109 genes) distributed among pseudochromosomes III, VI, VII, VIII, X, and XVIII. However, when orthologous gene pairs included disrupted genes and genes with more degenerated sequences, where AM560 CDSs with less than 50% sequence similarity and 50% sequence coverage were also lifted, gene synteny became lower and more inversions could be identified (Supplementary table 7).

We further validated and improved the transferred gene models using highly accurate transcript sequences generated via PacBio Iso-Seq (Supplementary table 8). Fifteen thousand annotated gene models could be validated, including 20,000 annotated transcripts and 4,000 novel transcripts from known genes. Two hundred of such novel transcripts represented fusion transcripts from over 100 genomic loci, each spanning two to three reference genes (Supplementary table 9, Figure 5d).

Interestingly, a small fraction of PacBio Iso-Seq transcripts overlapped with genomic loci of disrupted genes described above. For example, in the TME204 H2 assembly, 546 Iso-Seq transcripts overlapped with 355 disrupted genes, 50% of these Iso-Seq transcripts matched the lifted incomplete transcript models, the rest represented novel transcript variants (Supplementary figure 7). The lifted gene annotation was also improved with 400 novel genes and associated 4,000 novel transcripts (Supplementary table 9, Figure 5 b,c). GO enrichment analysis suggested that the novel genes were mainly related to molecular function (MF) of various lyase activity, and biological processes such as response to stress/stimulus, development of meristem, tissue, and anatomical structures. (Supplementary figure 8).

**Figure 7.**
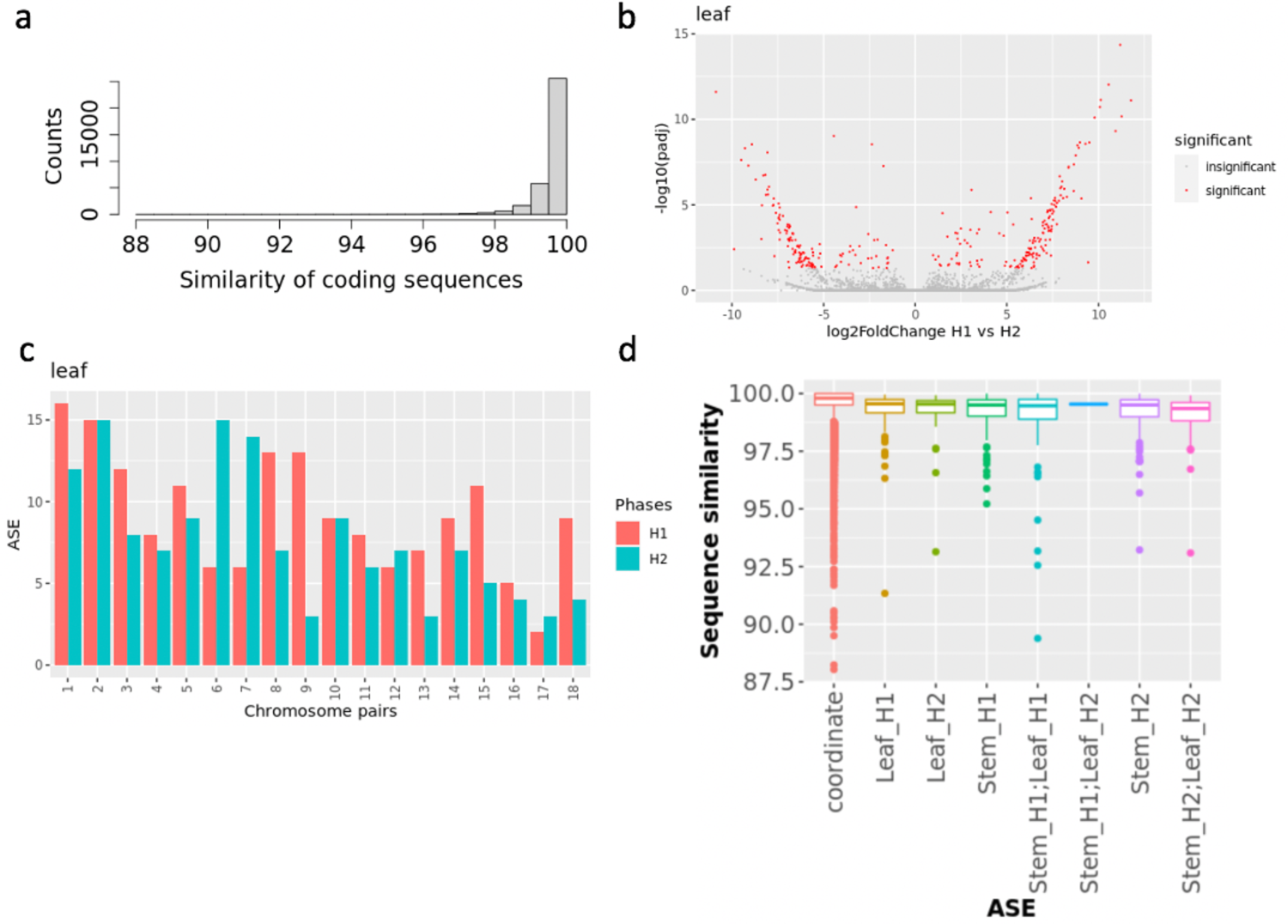
Allele-specific expression in cassava TME204. (a) Coding sequence similarity of the 34,194 bi-allelic cassava TME204 transcript pairs. (b) Identification of bi-allelic transcript pairs with significant allele-specific expression (ASE) differences. Red dots indicate genes with significant (adjusted p-value < 0.05) ASE differences, and gray dots represent genes without ASE differences. H1: TME204 haplotype 1 alleles; H2: TME204 haplotype 2 alleles. (c) Distribution of significant ASE alleles between pseudochromosome pairs in the TME204 diploid genome. (d) Classification of significant ASE alleles by ASE expression pattern in TME204 leaf and stem tissues.

**Figure 8.**
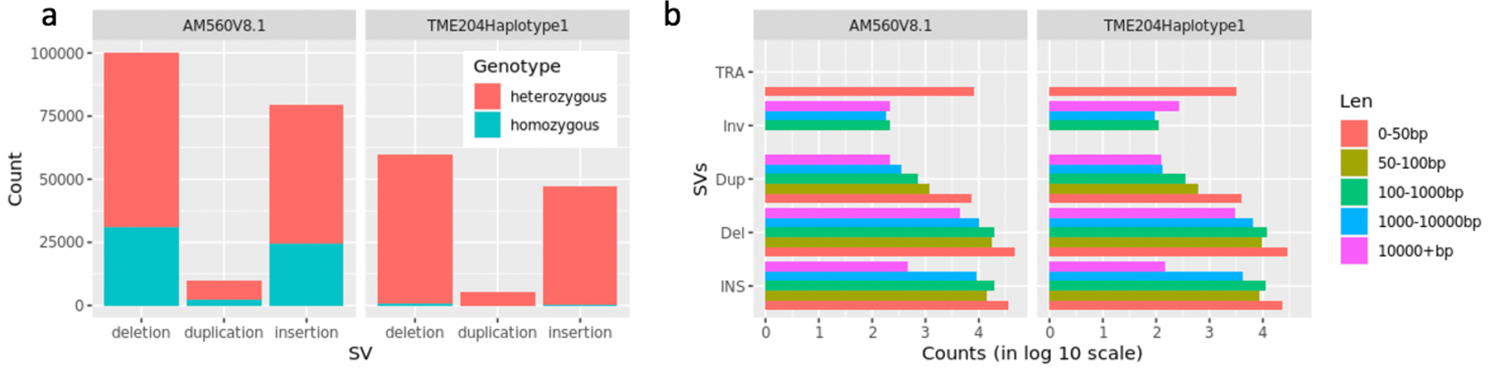
Identification of TME204 and AM560 structural variants (SVs) by reference guided analysis of HiFi read alignments. (a). Classification and counts of SVs by genotypes. (b) Classification and counts of SVs by variant types and length. INS: insertions; Del: deletions; Dup: duplications; INV: inversions; TRA: breakpoints of complex variants with unknow sizes, such as translocations etc.

### Tissue specific differentially expressed transcripts

Our haplotype-resolved genome assembly enabled annotations of 53,000 known transcripts and 8,000 novel transcripts in each haplotype assembly. Most transcripts have different sequences between haplotypes (Figure 6 a). In such cases, analyzing RNA-seq data using one haploid set of genes/transcripts as the reference could potentially miss haplotype-specific, novel expression patterns. Therefore we re-analyzed the previously published [31] RNA-seq data (Supplementary table 10) generated from two different tissues (TME204 leaf vs. stem) using our reference transcriptome of 119,805 unique transcripts, including both haplotype-specific isoforms and allelic pairs of common isoforms. In TME204 leaf and stem, 64,992 (54%) of transcripts were expressed, 6,696 (6%) transcripts showed significant difference in expression (Fold change above 4, adjusted p-value < 0.001). When compared among single-copy transcripts with multi-copy ones (i.e. same transcripts from duplicated gene loci), relatively lower fraction of multi-copy transcripts was expressed and differentially regulated (Figure 6 b). This is consistent with previous findings [32] that single copy genes are generally more highly expressed than multi-copy genes. When investigating the gene loci giving rise to tissue-specific differentially expressed transcripts (DET), we identified many gene loci where the associated DETs were solely from H1 or H2 (Figure 6 c,d). GO terms related to the cell wall macromolecular metabolism process were enriched in these haplotype-specific DETs, while GO terms related to the photosynthesis process were enriched in DETs that were coordinately regulated in both haplotypes, suggesting that these two groups of DETs play very different roles in cassava leaf and stem transcriptomes.

### Isoform allele-specific expression

For isoforms that are common to both haplotypes, we further investigated allele-specific expression (ASE) differences between the 34,194 bi-allelic transcripts from 25,636 orthologous gene loci (see Materials and Methods). Most of these bi-allelic pairs maintained high levels of coding sequence similarity (Figure 7 a). Only 2-3% of the alleles showed significant (adjusted p-value < 0.05) differences in expression between allelic pairs in leaf and stem, respectively. Most alleles were coordinately expressed in the TME204 genome. But alleles with significant expression differences showed mainly large fold changes (Figure 7 b) and on average shared slightly lower sequence similarity (Figure 7 d). The number of significant alleles between allelic chromosome pairs was similar for some chromosome pairs but varied greatly for the others (Figure 7 c), suggesting that distributions of ASE could be biased at some genomic loci between the haplotypes. In total, 119 alleles showed consistent expression biased towards one haplotype in both stem and leaf. There was only one transcript (Manes.05G108100.1, xyloglucan:xyloglucosyl transferase TCH4) that displayed switched allele expression in leaf vs. stem.

### Intra- and inter-genomic diversity of cassava genomes

Based on k-mer analysis, each TME204 haplotype harbors close to 20% of haplotype-specific k-mers. However, analysis of orthologous pairs of coding sequences revealed high gene synteny and coding sequence similarity on average. To systematically investigate sequence differences between the TME204 haplotypes and between cassava cultivars, we produced reliable alignments between assembled sequences longer than 500 bp, with exact matches >100 bp [33, 34]. In each haploid genome, a significant percentage of sequences was too divergent to be aligned between the TME204 paired haploid genomes (24%, 181 Mbp) or between the TME204 and AM560 cultivars (29%, 220 Mbp). The average level of sequence differences between aligned sequences from the two TME204 haploid genomes was 1.12%, including 2,526,852 SNPs and 1,733,059 single nucleotide indels (Table 4). Assemblytics analysis of these reliable alignments identified 13,332 small indels (20-50 bp) and 13,213 large indels (50 −10,000 bp) between the two TME204 haplotypes. Generation of the large indels seems to be repeat driven, 67% of which were expansion/contraction of repetitive elements, while only 3% of the small indels were of the same types (Table 4, Supplementary figure 9). The levels and characteristics of inter-genomic differences between the two cassava cultivars (TME204 vs. AM560) were similar to those within TME204 diploid genome (Table 4, Supplementary figure 9).

**Figure 9.**
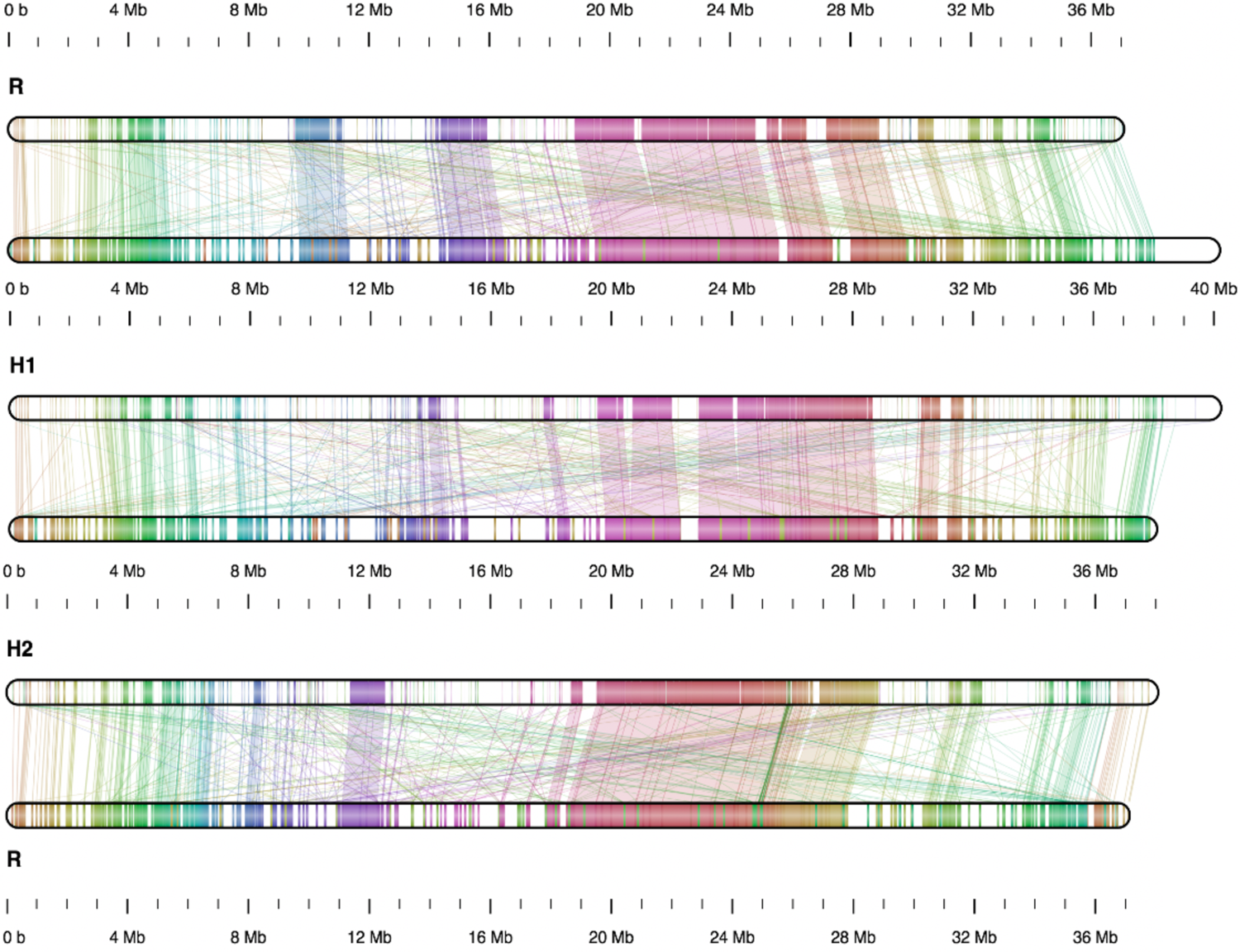
Chromosome XII maps of TME204 and AM560 show extensive genomic rearrangements between the chromosome pairs. “R” indicates the pseudochromosome from the reference AM560 v8.1 assembly, “H1” pseudochromosome from the TME204 haplotype 1 assembly, “H2” pseudochromosome from the TME204 haplotype 2 assembly. Shared regions between chromosome pairs are shown as color segments and connected by color lines between chromosomes. Shared regions with similar sequence information content were detected by Smash++ with parameters adjusted for highly repetitive genomes (Materials and Methods). White segments represent regions that are degenerated between a chromosome pair. Such accumulation of degenerated genomic sequences was observed between all pseudochromosome pairs, both within the TME204 diploid genome, and between each TME204 haplotype and the AM560 haploid genome (Supplementary File 5).

In our gene synteny analysis, we noticed that the number of inversions between any two cassava haploid genomes increased when regions with more degenerated sequences were included. To categorize inversions in a more comprehensive approach, we first compared HiFi reads directly to AM560 contigs and TME204 haplotigs, which identified indels, inversions and breakpoints of other complex SVs such as translocations, etc. If the TME204 genome was assembled error-free, all sequence variants between one TME204 haplotype and all HiFi reads would have been heterozygous and representing intra-genomic diversity. Indeed, only less than 1% of structural variants (SVs) reported by HiFi read alignments were homozygous. They could have resulted from mis-assemblies and/or mis-alignments. Most of the SVs (> 99%, 115,000) were heterozygous between TME204 haplotypes, confirming that our TME204 haplotigs are structurally accurate and do harbor a high level of intra-genomic sequence differences. The very high number of reported SVs was due to the high sensitivity of the analysis method, since SVs supported by three or more HiFi reads could be identified with high confidence. Similarly, between the TME204 diploid genome and the AM560 genome, 198,000 SVs were identified by the HiFi read alignments, of which 70.5% were heterozygous and specific to only one of the TME204 haplotypes (Figure 8 a). On average, the number of SVs between one TME204 haplotype and AM560 haploid genome reached 128,000, which is again very similar to the number of intra-genomic SVs (115,000) between TME204 haplotypes. Besides the much higher sensitivity, analysis of HiFi read alignments was also able to identify very small inversions such as those from 100 bp to a few Kbp (Figure 8b), which were not captured by gene synteny analysis. Consequently, the number of inversions reported with this method was much higher and not directly comparable with gene synteny analysis.

To tackle this limitation, we decided to investigate the TME204/AM560 pseudochromosome pairs by identifying and examining regions that shared information content [35], which is more robust in comparing sequences with low sequence identity and where the linear order of homologs is not preserved [36]. The analysis revealed that each cassava pseudochromosome consists of islands of conserved regions flanking by regions with more degenerated sequences. Although the order of these conserved regions was mostly kept between each pseudochromosome pair, extensive genomic rearrangements still exist (Figure 9). In total, more than 2,500 inversions were detected between each pair of cassava haploid genomes with this method (SupplementaryDataFile.pdf).

### Cassava pan-genome

The presence of haplotype-specific k-mers and abundant SVs between the cassava haploid assemblies suggests that any of the linear reference genomes of one haplotype, either the AM560 pseudo-haplotype or TME204 H1 or H2, cannot represent the sequence diversity of cassava populations and may miss haplotype-specific sequences. To overcome this limitation, we built a pan-genome graph from TME204 H1 and H2, and also one including the reference AM560 pseudo-haplotype. Starting with each initial reference haplotype (TME204 H1 or AM560), haplotype-specific large SVs (100 bp and 100 kbp) were identified in the query haplotype and subsequently amended to the reference haplotype for pan-genome graph reconstruction. We found 114,773,684 bases representing 40,776 such large SVs in TME204 H2 that were divergent from TME204 H1 (Figure 10 a). In comparison to the linear TME204 H1 as the only reference genome, using the TME204 pan-genome as reference allowed us to map more Illumina reads from the same TME204 sample with higher accuracy (i.e. mapping quality 20 and above) (Figure 10 b). In the pan-genome that includes the AM560 genome and the two TME204 haplotypes, we found 198,028,264 bases representing 53,098 large SVs in the two TME204 haplotypes that were divergent from AM560. As reported above by the Assemblytics analysis, where a majority of large indels (50 bp – 10 kbp) are expansion/contraction of repeats, the SV harboring divergent sequences in both pan-genomes are enriched for repeats, especially LTR elements (Figure 10 c), suggesting that most SVs captured by pan-genome graphs are LTR retrotransposons related.

**Figure 10.**
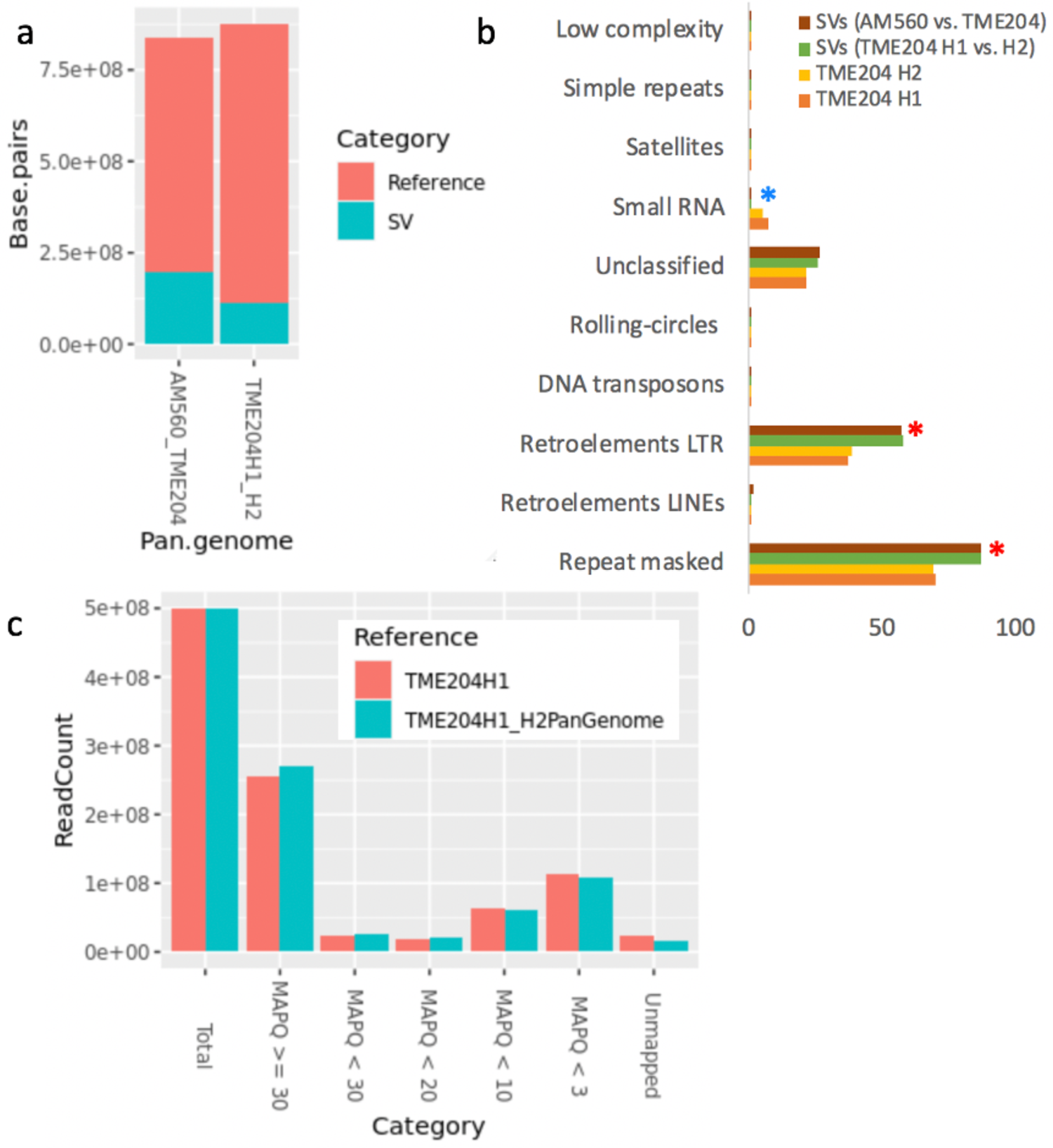
Properties of cassava pan-genomes. (a) Cassava pan-genomes across different assemblies and large SVs (100 bp – 100 kbp) detected by pan-genome graphs. The pan-genome size decreased when AM560 was included because there were less 1-to-1 orthogonal regions between AM560, TME204 H1, and H2. (b) SVs in cassava pan-genomes are enriched with repeats (Chi square test p-vale < 0.05), especially LTR elements (p-value < 0.05), and are deprived of small RNAs (p-value < 0.005). (c) The pan-genome of TME204 H1 and H2 improved mapping rate and mapping quality of Illumina PE reads collected from the same DNA sample.

## Discussion

By comparing PacBio CLR and HiFi sequencing technologies and benchmarking four HiFi assemblers [14,18,22], we demonstrate that HiFi reads are extremely effective in producing a nearly complete and accurate haplotype-resolved assembly of the complex diploid cassava genome. The combination of high base accuracy and long read length greatly simplified the data analysis workflow, decreased data footprints, shortened data analysis time, and improved the assembly quality. CLR-Falcon assembly starts with read self-correction, which is not only computationally expensive, but can also mix reads from different haplotype alleles, paralogous gene members, or repetitive elements. In contrast, HiFi reads have higher resolution and accuracy in resolving these sequence variants. All HiFi TME204 assemblies reached consensus accuracy between Q40 (99.99%) and Q50 (99.999%). The CLR-Falcon contig sequences were less accurate even after extensive polishing using signal level data, which also has the risk of introducing novel errors because current polishing pipelines cannot accurately differentiate reads from different haplotype alleles and repeat copies [23].

Among the compared HiFi assemblers, hifiasm generated the most completely haplotype-resolved TME204 genome assembly. The haplotigs reached NG50 of 18 Mbp, with consensus accuracy of QV45. Using Hi-C technology and the cassava genetic map, we reconstructed the TME204 diploid genome into 18 pairs of pseudochromosomes, with three pairs as haplotigs without sequencing gaps. These values satisfy the 6.7.Q40 and 7.C.Q50 genome assembly quality metrics, which are measures for close-to-finished genome qualities as proposed by the VGP consortium [23], further emphasizing the high completeness and quality of the assembled TME204 genome.

The sequencing strategy of HiFi in combination with Hi-C not only enabled the assembly of haplotype resolved chromosome pairs, but also allowed reconstruction of over 300 mitochondrial haplotigs with lengths varying between 25 to 76 kbp. Plant mitochondrial genomes are known to be highly fragmented, with total lengths varied from 200 to 2,000 Kbp [37]. The 53 mitochondrial haplotigs in the TME204 H2 assembly added up to a total size of 2 Mbp (Table 3), which can represent a complete mitochondrial genome. Interestingly, there were still 281 mitochondrial haplotigs (with a total length of 12 Mbp) in the TME204 H1 assembly, suggesting the presence of different sequence variants of the mitochondrial genome. This result strongly supports the recent discovery of plant mitochondrial genomes as a complex and dynamic mixture of sequence variants (37). It signifies that the highly accurate base information over very long stretches of DNA molecules provided by the combination of HiFi sequencing with Hi-C technology is powerful in resolving the complexity of multiple haplotypes and isoforms, which will revolutionize and fundamentally improve future assemblies of plant genomes.

Annotation of the haplotype resolved TME204 genome using reference gene models and Iso-Seq transcripts revealed a complex and dynamic transcriptome of cassava. We found that different sets of known transcripts have become disrupted in the two haploid genomes, potentially leading to changes in biological functionality. Among the genes lifted from AM560, 9% appeared as disrupted genes in TME204 haplotypes. However, expression of some of these disrupted genes was supported by Iso-Seq transcripts, similar to the expression of fusion genes. In humans, tissue-specific pseudogene expression has been reported [39] and fusion transcripts are found to play an important role in tumorigenesis [40]. Such complex transcripts and their functions in plants are underexamined but efforts have been made to start the exploration [41, 42]. It requires further investigation to determine if the expressed fused or disrupted genes we found in TME204 are real or result from annotation artifacts.

The haplotype resolved TME204 transcriptome revealed that a majority of the transcript sequences are not identical between the two haplotypes. The reference transcriptome representing all isoforms enabled us to identify haplotype-specific isoforms that were differentially expressed in different tissue types. For common isoforms shared between haplotypes, most alleles are coordinately expressed, as recently reported in ginger [43]. For isoforms showing ASE, the expression bias is either tissue-specific or retained in different tissues. Only one transcript switched the expressed allele between leaf and stem tissues. This is also similar to the patterns observed in ginger and tea plant [43, 44]. We previously reported more genes with ASE for cassava 60444 and TME3 [8] because different RNA isoforms (i.e. different exon usage) also contributed to expression differences between orthologous gene pairs. These different RNA isoforms were excluded in the current ASE analysis of isoform alleles.

Each current TME204 H1 and H2 assembly is still a random mixture of different parental chromosomes because with Hi-C technology alone it is not possible to phase across chromosomes. Trio-binning [45] using two parental genomes will be needed to completely separate parental chromosomes in the offspring genome and to assist in the analysis of monoallelic expression of parentally imprinted genes in offspring. The findings of haplotype-specific disrupted genes, haplotype-specific DETs and ASE RNA isoforms will still hold true after reshuffling of pseudochromosomes between haplotypes, although the actual functional GO terms enriched within the specific set of incomplete transcripts/DETs/ASE isoforms may change. Our haplotype resolved transcriptome will be a powerful resource and tool for establishing new technologies, such as novel marker identifications and genome editing for cassava trait improvement and breeding.

Extensive SVs and divergent sequences per haploid genome are dispersed throughout both haplotypes, and the levels of intra-genomic and inter-genomic diversity are similar in cassava. Genome regions with SVs are enriched with repeats, especially LTR elements. Accumulation of SVs and hemizygous sequences have been recently reported for other crops such as grapes, potatoes, and rice, and are considered a major force contributing to the cost of domestication [19,46,47].

Analysis of SVs in cassava TME204 population samples will help to reveal to what extent SV is driving cassava genome evolution. Our study demonstrates that reference-guided analysis of HiFi read alignment is more sensitive in identifying SVs than comparative analysis of assembled consensus sequences, which will be a cost-effective method for population scale analysis of SVs.

The high degree of genomic variations in cassava cultivars also highlights the importance of building a pan-genome[48–51] for research and breeding. Under-representation of genetic diversity by any linear haploid cassava genome will limit our understanding of genetic variations in reference-guided analysis, especially when samples are sequenced using Illumina short reads, for example in genotyping-by-sequencing and RNA-seq experiments. Haplotype-specific short reads may remain unmapped, thus important genome information may be left undiscovered. Technically, large SVs are a frequent source of errors in aligning short Illumina reads, which may lead to mis-interpretation of data <u>(1–4)</u>. We demonstrate that using a pan-genome reference did increase mapping rate and mapping quality of Illumina reads in comparison to using a conventional linear haploid reference. Detailed investigation of a cassava pan-genome, including more cultivars, and its influence on interpretations of omics data is on-going and will be reported in the near future.

### Potential implications

Using the HiFi sequencing strategy in combination with Hi-C, we reconstructed two chromosome scale haploid genomes for the diploid cassava TME204, which allowed us to study the sequence, gene content, gene expression, and genome structure with unprecedented resolution. The haplotype resolved genome and transcriptome will be a valuable resource for cassava breeding and research. The ability to resolve the high complexity of multiple haplotypes and isoforms demonstrated in our study will provide insights for future work on plant genomics.

## Methods

### DNA extraction and Illumina shotgun sequencing

Leaves were collected from 6- to 8-week old *in vitro*-grown TME204 plants. Genomic DNA was extracted using DNeasy Plant Mini Kit (QIAGEN). The TruSeq DNA Nano Sample Prep Kit v2 (Illumina) was used in subsequent steps of library preparations. The DNA sample (100 ng) was sonicated with the ME220 Focused-ultrasonicator (PN: 500506, Covaris) using settings specific to the fragment size of 350 bp. The fragmented DNA sample was size-selected using AMpure beads (Beckman Coulter), end-repaired and adenylated. TruSeq adapters containing Unique Dual Indices (UDI) (GACACCATGT and GCACGGTACC) for multiplexing were ligated to the size-selected DNA sample. Fragments containing TruSeq adapters on both ends were selectively enriched by PCR. The quality and quantity of the enriched library were validated using Tapestation (Agilent Technologies). The product is a DNA fragment population with an average fragment size of 500 bp. The library was adjusted to 10nM using a Tris-Cl 10 mM, pH8.5 with 0.1% Tween 20 buffer. The Novaseq 6000 (Illumina) was used for cluster generation and sequencing according to the standard protocol for paired-end (PE) sequencing at 2 X150 bp.

### High molecular weight DNA extraction

Fresh leaves were harvested from *in vitro*-grown TME204 plants kept in the dark for 12-24 hours pre-harvest, and the petiole and basal midrib were removed with a sterile pair of scissors. One gram of leaf tissue was then snap-frozen in liquid nitrogen and homogenized to a powder with a mortar and pestle. Lysis buffer (9.5 mL of G2 buffer from the Blood & Cell Culture DNA Midi Kit (QIAGEN) and 19 µL of RNase A (100 mg/mL, Sigma Aldrich) was added to the homogenized tissue in a 50 mL conical centrifuge tube (Falcon). 500 µL of Proteinase K (20 mg/mL, Roche) was then added to the sample and the mixture was vortexed for 10 seconds. The sample was incubated at 50°C (Memmert Incubator) on a lab roller for 3 hours. Afterwards the sample was centrifuged for 10 minutes at 20°C at 1,800 x g. The supernatant was then used for high molecular weight (HMW) genomic DNA extraction according to the Genomic-tips protocol (100/G, Blood & Cell Culture DNA Midi Kit, QIAGEN).

### PacBio CLR and HiFi library preparation and sequencing

The PacBio CLR and HiFi libraries were produced using the SMRTbell Express Template Prep Kit 2.0 (Pacific Biosciences) according to the manufacturer’s instructions. The concentration of HMW genomic DNA was measured using a Qubit Fluorometer dsDNA Broad Range assay (Thermo Fisher Scientific). The CLR and HiFi library preparations started with 8 μg and 15 μg HMW DNA, respectively. The DNA samples were mechanically sheared to an average size distribution of 30 kbp (CLR) and 20 kbp (HiFi) using a Megaruptor Device (Diagenode). A Femto Pulse gDNA analysis assay (Agilent Technologies) was used to assess the fragment size distribution. Sheared DNA samples were DNA damage-repaired and end-repaired using polishing enzymes. PacBio sequencing adapters were ligated to the DNA template. For the CLR library, a Blue Pippin device (Sage Science) was used to size select DNA fragments > 25 kbp. For the HiFi library, a Sage Elf device (Sage Science) was used to enrich DNA fragments > 15 kbp. The size selected DNA libraries were quality-checked and quantified using a Femto Pulse gDNA analysis assay and a Qubit Fluorometer, respectively. The CLR SMRT bell template-polymerase complex was created using the Sequel binding kit 3.0 (Pacific Biosciences) and subsequently sequenced on a PacBio Sequel instrument using the Sequel Sequencing Kit 3.0 (Pacific Biosciences) with six Sequel™ SMRT® Cells 1M v3 (Pacific Biosciences), taking a 10-hour movie per cell. The HiFi SMRT bell template-polymerase complex was created using the Sequel II Binding Kit 2.0 and Internal Control 1.0 (Pacific Biosciences), sequenced on a PacBio Sequel II instrument using the Sequel II Sequencing Kit 2.0 (Pacific Biosciences) and one Sequel™ II SMRT Cell 8M (Pacific Biosciences), taking a 30-hour movie.

### Hi-C library preparation and sequencing

Two grams of fresh leaf tissue was harvested from *in vitro*-grown TME204 plants and flash-frozen in liquid nitrogen. The leaf tissue was then shipped in dry ice to Arima Genomics (San Diego, USA) for Hi-C library preparation and sequencing. Flash-frozen leaves were first crosslinked, followed by Hi-C library generation using the High Coverage Arima Hi-C kit (PN: A410110). Illumina-compatible sequencing libraries were prepared by first shearing purified proximally-ligated DNA and then size-selecting DNA fragments using SPRI beads (Beckman Coulter). The size-selected fragments containing ligation junctions were enriched using Enrichment Beads provided in the High Coverage Arima Hi-C kit and converted into Illumina-compatible sequencing libraries using the Swift Accel-NGS 2S Plus DNA Library Kit (PN: 21096). After adapter ligation, DNA was PCR-amplified and purified using SPRI beads. The purified DNA was quality-controlled using qPCR (Roche) and Bioanalyzer (Agilent Technologies), then sequenced on the Illumina HiSeq X following manufacturer’s protocols, yielding 727,211,240 read pairs (2X150 bp) (Accession number: ERR5484651).

### RNA isolation, PacBio Iso-Seq library preparation and sequencing

Three different tissues were collected from greenhouse-grown TME204 plants: the top five leaves with petioles, apical and lateral meristems including the stem, and fibrous roots. The various tissues were flash-frozen in liquid nitrogen and homogenized with a mortar and pestle. RNA was isolated with the Spectrum Plant Total RNA kit (Sigma-Aldrich) according to Protocol A. The quantity and quality of total RNA samples were measured using Qubit RNA BR Assay Kit (Thermo Fisher Scientific) and Agilent TapeStation 4200 with RNA-specific tapes (Agilent Technologies), respectively. Samples with RNA integrity numbers ≥ 7 were used for Iso-Seq library preparation and sequencing.

PacBio Iso-Seq templates were prepared using the NEBNext Single Cell/Low Input cDNA Synthesis & Amplification Module (New England BioLabs) and PacBio Iso-Seq Express Template Switching Oligos (TSO) (Pacific Biosciences), following the PacBio Iso-Seq protocol “Procedure & Checklist – Iso-Seq Express Template Preparation for Sequel and Sequel II Systems” (PN 101-763-800). In detail, RNA samples (300 ng) were reverse-transcribed with oligo-dT primer in combination with the 5’ template-switching oligonucleotide (TSO). Synthesized first-strand cDNAs from different tissues were multiplexed in the cDNA amplification reaction with barcoded forward and reverse cDNA PCR primers annealing to the sequences of the 5’ TSO and 3’ oligo-dT primer. Barcoded and amplified cDNAs were purified using ProNex beads (Promega), following the workflow targeting transcripts around 2 kb in length. Purified ds cDNAs were used for PacBio template preparation with the SMRTbell Express Template Prep Kit 2.0 (PN: 100-938-900) (Pacific Biosciences). Afterwards the Iso-seq SMRT bell template-polymerase complex was prepared using Sequel II Binding Kit 2.1 (Pacific Biosciences) and PacBio sequencing primer v4, and was subsequently sequenced on a PacBio Sequel II instrument using Sequel II Sequencing Kit 2.0 (Pacific Biosciences) and single Sequel™ II SMRT Cell 8M (Pacific Biosciences) taking a 30-hour movie.

### Bacterial artificial chromosome (BAC) clone library construction, screening, sequencing and assembly

High molecular weight (HMW) DNA was prepared from TME204 young leaves as previously described [53, 54]. Agarose embedded HMW DNA was partially digested with HindIII (New England Biolabs), sized through two size selection steps by pulsed field gel electrophoresis (CHEF Mapper system, Bio-Rad Laboratories) and ligated into the pAGIBAC-5 HindIII-Cloning vector. Pulsed-field migration programs, electrophoresis buffer and ligation desalting conditions were done according to [55]. The insert size of the BAC clones was assessed using the FastNot I restriction enzyme (New England Biolabs) and analyzed by pulsed field gel electrophoresis. Colony picking was carried out using a robotic workstation QPix2 XT (Molecular Devices) using a white/blue selection. White colonies were arranged in 144 384-well (55,296 BAC clones) microtiter plates containing LB medium with chloramphenicol (12.5 μg/mL) supplemented with 6% (v/v) glycerol.

Individual BAC clones were selected using radiolabeled ([α-33P]dCTP) probes. DNA were extracted from individual clone using Nucleobond Xtra midi kit (Macherey-Nagel) and used for PacBio library preparation by The French Plant Genomic Resources Center (CNRGV) of the French National Research Institute for Agriculture, food and Environment (INRAE). PacBio sequencing was performed on the Sequel II system with a movie time of 30 hours with 120 min pre-extension step by Gentyane Genomic Platform (INRAE). Circular consensus sequence (CCS) reads per BAC clone were generated using SMRT Link (v9.0.0), and assembled using hifiasm (v0.12.0). More details on BAC clone screening, sequencing and assembly can be found in supplementary materials and methods.

### Sequencing data quality control

The technical quality and potential sample contamination in Illumina PE reads were evaluated using FastQC (v 0.11.8) (https://www.bioinformatics.babraham.ac.uk/projects/fastqc/) and FastqScreen (v 0.11.1) ( https://www.bioinformatics.babraham.ac.uk/projects/fastq_screen/), respectively. The technical quality of PacBio raw data was checked using the “QC module” in the PacBio SMRT Link software (version 8.0) (https://www.pacb.com/support/software-downloads/). The technical quality of Hi-C data was checked using HiCUP (v0.8.0)[56].

### Estimation of genome properties

Estimation of genome complexities such as repeat content and the level of heterozygosity were made with k-mers in the Illumina PE reads using Preqc in SGA (v 0.10.15) [57, 58]. Analyzed datasets and their accessions are: Human (ERR091571-ERR091574) [58], cassava AM560 (SRR2847385), Cassava TME204 (ERR5484652), cassava 60444 (ERR5484654) (8), cassava TME3 (ERR5484653) (8).

### PacBio CLR and HiFi whole genome assembly

PacBio CLR reads were assembled using Falcon [22] in pb-assembly (v0.06). PacBio HiFi reads were assembled using multiple HiFi specific assemblers, including Falcon in pb-assembly (v0.0.8), Improved Phased Assembler IPA (v1.0.5) (https://github.com/PacificBiosciences/pbipa), hifiasm (v0.7) [18], and HiCanu (v2.0) [14]. Default options were used unless otherwise noted. Improved phased assembly (IPA) was run with both phasing and polishing included.

### Benchmarking analysis of assembly accuracy and completeness

Assembly statistics were collected using QUAST (v4.5) [59]. NG50 [26] was calculated using the haploid genome size of 750 Mbp. Base-level accuracy and completeness was measured using both mapping-based and k-mer-based methods. TME204 Illumina PE reads were mapped to all genome drafts using BWA mem (v0.7.17) [60]. Statistics of read mapping were collected using samtools (version 1.10) [61] and Qualimap (v2.2.1) [62]. Sequence differences between mapped reads and assembled sequences, and fractions of mapped Illumina PE reads were used to measure consensus accuracy and assembly completeness. Accuracy and completeness of all drafts were further estimated from the k-mers present only in assembled sequences and recovery rate of reliable Illumina k-mers (v1.1) [27]. Briefly, Meryl (v1.7) was used to identify all k-mers with k=20 present in TME204 Illumina PE reads. The k-mer size of 20 was selected based on a haploid genome size of 750 MB and a diploid genome size of 1.5 Gbp. In Merqury, k-mers in each assembly were evaluated for their presence in the Illumina k-mer spectrum. A k-mer missing in the Illumina set is counted as a base-level ‘error’. The fraction of such ‘erroneous’ k-mers was used to calculate a phred scale consensus accuracy quality value (QV). A QV score of 40 means 1 in 10,000 k-mers was specific to the assembled sequences and missing from the Illumina reads. Assembly completeness was measured by k-mer completeness, which is the fraction of reliable Illumina k-mers retained in the assembly (v1.1).

For evaluation of structural accuracy, Merqury k-mer analysis results were first used to compute false duplication rates, where k-mers that appeared more than twice in each haploid assembly were used to identify artificial duplications. PacBio CLR reads were then aligned to each haploid genome and the coverage was analyzed using Asset software (https://github.com/dfguan/asset). Assembled regions supported by 10 and more PacBio CLR reads were identified as reliable regions.

Functional completeness was measured using BUSCO (v5) completeness of single-copy orthologs discovered in plants (Viridiplantae Odb10) [63], and alignment rates of reference genes and TME204 Iso-Seq transcripts. The AM560 reference genome (v8.0) and gene annotation (v8.1, https://phytozome-next.jgi.doe.gov/info/Mesculenta_v8_1) [30] were downloaded from JGI Phytozome 13 (https://phytozome-next.jgi.doe.gov/) [64]. Reference coding sequences (CDSs) were aligned to the AM560 reference genome and TME204 haplotigs using minimap2 (v2.15r905, - cxsplice -C5) [65]. “asmgene” completeness and duplication scores [18] were calculated using the “paftools” script from the minimap2 package, based on CDSs mapped at ≥97% identity over ≥99% of the CDS length. Iso-Seq data collected from TME204 transcriptomes of fibrous root, stem meristems and leaves were spliced aligned using minimap2 (v2.15r905, x splice:hq). Alignment statistics were collected using alignqc [66].

### Haplotype-resolved, phased contig assembly using HiFi reads integrated with Hi-C technology

Two sets of haplotype-resolved, phased contig (haplotig) assemblies were generated using hifiasm (v0.15.3) with a combination of HiFi reads and paired-end Hi-C reads. Haplotigs were first validated against the high dense genetic map of cassava [29], which contains 22,403 SNP markers with allele numbers ranging from 2 to 6. Allelic sequences (50 nt upstream sequence + allele sequence + 50 nt downstream sequence) were aligned to haplotigs using BLAT (v3.2.1)[67]. For each haplotig, correlation plots of genetic vs. physical distance based on uniquely and perfectly aligned alleles were generated for visual inspection. Sequences of BACs were also aligned to haplotigs using BLAT (v3.2.1). The best BAC-to-haplotig alignment was manually inspected to identify resolved BACs, where one continuous BAC-to-haplotig alignment was produced.

### Construction of Pseudochromosomes

Hi-C reads were mapped back to each set of haplotigs independently using the Arima mapping pipeline (https://github.com/ArimaGenomics/mapping_pipeline) and were used to further scaffold haplotigs with SALSA2 (v2.2, assisted by the assembly graph, resolved mis-assemblies, five iterations) [68]. No haplotigs in TME204 H1 were further scaffolded with Hi-C data after five rounds of iteration (Supplementary file 1). Thirty haplotigs in TME204 H2 were scaffolded into 13 scaffolds, of which seven were chromosome-scale and consistent with the genetic map (Supplementary table 5). One scaffold was apparently a technical artifact based on genetic markers, reaching 107 Mbp long and joining haplotigs together from chromosomes I, XVI and XVIII (Supplementary table 5).

Chromosomes VII, IX and XI were not reconstructed in H2. Because Hi-C scaffolding did not generate results for H1 and the results for H2 were not all satisfactory, ALLMAPS (v0.8.12) [68] was used to reconstruct pseudochromosomes for both sets of haplotigs based on the genetic map [29]. Given the observed high congruence between the map and haplotigs, as well as between the map and Hi-C scaffolds, our choice of scaffolding strategy was reasonable and sound.

### Repeat modeling, genome masking and annotation

Starting with the assembly of all resolved alleles (i.e. primary plus alternate contigs), repeat elements were predicted using RepeatModeler (v2.0.1), with dependency on TRF (v4.09) [69], RECON (1.08) [70], RepeatScout (v1.0.6) [71], and RepeatMasker (v4.1.0) [72]. Analysis of Long Terminal Repeats (LTRs) were enabled with GenomeTools (v1.5.9) [73], LTR_Retriever (v2.9.0) [74], Ninja (v0.95-cluster_only) [75], MAFFT (v7.471-with-extensions) [76] and CD-HIT (v4.8.1) [77]. Among the 1436 predicted repeat families, 1021 were unknown/novel according to RepeatClassifier (V2.0.1) [78]. Five predicted repeat families with significant hits to plant genes were identified and removed using ProtExcluder (v1.1) (http://www.hrt.msu.edu/uploads/535/78637/ProtExcluder1.2.tar.gz). Each TME204 haplotype assembly was then masked with the customized repeat library using RepeatMasker (v4.1.0).

Genome annotation was performed by first transferring reference gene models from AM560 to TME204 haplotype assemblies using liftoff (v1.6.1) [79]. Transferred gene models were further improved by adding novel transcripts and gene models based on TME204 PacBio Iso-Seq and Illumina RNA-seq data. In detail, Liftoff aligns AM560 genes and transcripts to each TME204 haplotype assembly and finds the mapping that maximizes sequence similarity while preserving the structure of exon, transcript, and gene. Analyzing sequence alignments at all levels of annotated features allows genes and transcripts to be lifted despite the presence of abundant SVs between the cassava genomes. Finding extra copies of the same genes was enabled with a minimum sequence identity of 95% in exons/CDSs. Synteny analysis of orthologous pairs of transferred genes was performed using MCScanX [80] and visually inspected using accusyn (https://accusyn.usask.ca/).

PacBio Iso-Seq reads from the three different TME204 tissue types were clustered into high quality (accuracy 99.9%, HQ) transcripts using the Iso-Seq Analysis Application in PacBio SMRT Link software (10.1.0.119588). HQ transcripts from different tissues were pooled and aligned to each haplotype assembly. Redundant isoforms were collapsed into 36,000 unique transcripts using cDNA_Cupcake (https://github.com/Magdoll/cDNA_Cupcake/). Unique transcripts were quality-controlled, filtered and classified using SQANTI3 (v4.1) [81] to identify novel transcripts and gene models. In detail, previously published Illumina RNA-seq data (Supplementary table 10) from TME204 [82] were used to check PacBio transcript coverage. Transcripts in which junction sites had low Illumina RNA-seq read support (<4) were filtered out. Transcripts showing other technical artifacts, such as polyA intra-priming when aligning to the haplotype assembly, were also excluded, yielding 29,000 accurate isoform sequences. They were compared against lifted reference gene models and transcripts for reference model validation and identification of novel transcripts and genes. Novel protein sequences were functionally annotated using interproscan (v5.52-86.0) [83].

### Differential expression of transcripts and analysis of allele-specific expression

Transcript sequences annotated in TME204 H1 and H2 were pooled and de-duplicated using cd-hit-est (v4.8.1) to generate a single set of reference transcripts for expression quantification and differential analysis of tissue. Transcripts duplicated between TME204 H1 and H2 were counted as homozygous alleles. Transcripts duplicated within a haploid genome were counted as multi-copy alleles. Duplicated transcripts were represented by a single reference transcript. RNA-seq reads previously generated from leaf and stem with three biological replicates (30) were mapped to the reference transcripts (119,805) representing two haploid transcriptomes using kallisto (v0.46.1). If a transcript expression value of at least two samples among the six given samples exceeded 1 TPM (transcript per million), we considered the transcript to be expressed. Differentially expressed transcripts (DETs) between tissue types were identified using DESeq2 (v1.32.0) as those with fold-change of TPM values between two tissue types greater than 4 and adjusted p-value <0.001.

For allele-specific expression (ASE), bi-allelic transcripts were identified by a combination of reciprocal blastn comparison of H1 and H2 transcripts and gene synteny. Among all transcripts associated with an orthologous gene pair, transcripts of reciprocal blastn best hits were considered as allele A and B. Expression values for bi-allelic transcripts were subset from the master quantification table including all resolved alleles, ASE was determined using the same package DEseq2, with adjusted p-value <0.05.

### Comparative genomics

For alignment-based sequence similarity analysis, the cassava reference genome AM560 was first disassembled into contig sequences using the utility function “split_scaffold” in IDBA (v1.1.3) [84]. Each set of TME204 haplotigs was then aligned to the AM560 reference contigs and against each other using nucmer (--maxmatch −l 100 -c 500) in MUMmer (v 4.0.0beta2) [34], which reported all sequence alignments longer than 500 bp with each exact match longer than 100 bp. Contigs rather than pseudochromosomes were used to prevent false positives when the padding Ns in the query did not match perfectly to the distance in the reference. Sequence alignments were further analyzed using dnadiff in MUMmer and Assemblytics [33] for identification of SNPs, single nucleotide indels, and large indels (20 bp - 10 kbp).

For chromosome level comparisons, the alignment free method smash++ (v20.04) [35] was used to identify similar/shared regions and genomics rearrangements larger than 10 kbp between pseudochromosome pairs. Parameters adjusted for analyzing highly repetitive genomes were: filter scale = large, filter size = 50000, filter type = blackman, threshold = 1.0, minimum segment size = 10000.

### Structural variant analysis using HiFi reads

TME204 HiFi reads were aligned to reference contigs of AM560 and TME204 haplotigs using minimap2 (v 2.15r905). SVs were called using PacBio structural variant calling and analysis tools (PBSV, https://github.com/PacificBiosciences/pbsv). Summary statistics of SVs were collected using SURVIVOR (v1.0.7).

### Pan-genome analysis

Pan-genomes were constructed using minigraph (v 0.15-r426) [52]. Large SVs (100 bp −100 kbp) were identified and extracted from each pan-genome graph using gfatools (0.4-r214) (https://github.com/lh3/gfatools).

### Gene ontology (GO) enrichment analysis

For each selected gene set, GO enrichment analysis was performed using topGO (v2.44.0) (https://bioconductor.org/packages/release/bioc/html/topGO.html) with Fisher exact test P-value cutoff set to 0.00001. Go annotations of the reference gene models were used as the background gene set, except for analysis of novel gene models, where GO annotations of novel gene models were also added as the background.

## Supporting information

All supplementary text, figures, and tables

## Declarations

## Ethics approval and consent to participate

Not applicable.

## Data availability

Raw sequencing reads from PacBio (HiFi, CLR and Iso-Seq) and Illumina (Hi-C and shotgun) were deposited in the European Nucleotide Archive (ENA) database under the accession number PRJEB43673 (or ERP127652 as the secondary accession number in ENA). Assembled genome sequences of TME204 H1 and H2 were deposited in the NCBI database under the accession number PRJNA758616 and PRJNA758615, respectively. Assembled BAC clone sequences were deposited in the NCBI GenBank database under the accession numbers MZ959795, MZ959796, MZ959797, and MZ959798. The five supplementary files and the version of annotation files used in the current analysis were uploaded to the Mendeley database (http://dx.doi.org/10.17632/fr6g4tgnfh.1#folder-dbb00a94-9bc5-4dad-a2bc-8da65fe270a0).

## List of abbreviations

ASE: allele-specific expression

BAC: bacterial artificial chromosome

BP: biological process

CCS: circular consensus sequence

CDS: coding sequence

CLR: continuous long reads

CMD: Cassava Mosaic Diseases

DE: differentially expressed/differential expression

DET: differentially expressed transcript

ENA: European Nucleotide Archive

GO: gene ontology

HiFi: high-fidelity

HMW: high molecular weight

Indel: insertion and deletion

IPA: improved Phased Assembler

MF: molecular function

NCBI: National Center for Biotechnology Information

numt’s: nuclear mitochondrial pseudogene regions

PacBio: Pacific Biosciences

PE: paired-end

QV: quality value

SMRT: Single Molecule Real-Time

SNP: single nucleotide polymorphism

SV: structural variation

TPM: transcript per million

UDI: Unique Dual Indices

VGP: the Vertebrate Genome Project

## Consent for publication

The cassava TME204 (Tropical Manihot esculenta 204) cultivar used in our study was obtained by ETH Zurich from the International Institute of Tropical Agriculture (IITA) in Nigeria in 2003 prior to the implementation of the International Treaty on Plant Genetic Resources for Food and Agriculture (https://www.fao.org/3/i0510e/i0510e.pdf). TME204 has been part of the ETH Zurich cassava germplasm collection since 2003. As a major crop, non-genetically modified cassava, including the wild type TME204 cultivar, is exempt from the Cartagena Protocol on Biosafety to the Convention on Biological Diversity (https://www.cbd.int/doc/legal/cartagena-protocol-en.pdf). The study reported in our manuscript follows all Swiss and international guidelines and legislation.

## Authors’ Contributions

WQ, YL, AP, RS and WG designed the study. YL and CC prepared DNA and RNA samples for sequencing. AP, SG and AB prepared CLR, HiFi and Iso-Seq libraries and performed PacBio sequencing. YL, NR, EP, SV and MF generated the BAC sequences. WQ, YL, PS, DP and WG analyzed data. WQ, YL, AP, AB, PS, and WG wrote the manuscript. All authors have reviewed the final manuscript before submission and have no competing interests.

## Acknowledgments

We thank the high-throughput sequencing team at FGCZ for Illumina sequencing service, Arima Genomics for Hi-C service, Dr. Daivd Stucki and Deborah Moine from PacBio for their technical support, Dr. Haoyu Chen from Harvard Medical School and Alaina Shumate from the Johns Hopkins University School of Medicine for insightful discussion. We thank Jay Tracy from FGCZ for reviewing the manuscript for English writing and clarity.

