## Supplementary material for "The haplotype-resolved chromosome pairs and transcriptome of a heterozygous diploid African cassava cultivar": All supplementary text, figures, and tables

**Supplementary material supporting the manuscript titled "Haplotype-resolved genome and transcriptome analyses of a heterozygous diploid African cassava cultivar"**

### Supplementary Methods

#### Screening of the BAC library

Radiolabeled probes ([α-33P]dCTP) were designed to screen the genomic BAC library. DNA fragments were PCR-amplified from TME204 genomic DNA using specific primers (Supplementary table 11).

High-density colony filters were prepared using a robotic workstation QPix2 XT (Molecular Devices, San José, CA, USA). BAC clones were spotted in duplicate using a 7 × 7 pattern onto 22 × 22 cm Membranes Hybond-XL filters (GE-Healthcare, Chicago, IL, USA). The whole BAC library was represented on one filter, containing 55 296 BAC Clones. After incubation at 37°C for 17 h, DNA fixation was processed as follows:

1. Denaturation on Whatman paper soaked with a solution of 0.5 m NaOH and 1.5 m NaCl for 4 min at room temperature and for 10 min at 100°C.
2. Neutralization on a Whatman paper soaked with 1 m Tris–HCl pH 7.4 and 1.5 m NaCl for 10 min, incubation in a solution of 0.25 mg ml−1 proteinase K (Sigma-Aldrich, St. Louis, MO, USA) for 45 min at 37°C and baking for 45 min at 80°C.
3. UV fixation on a Biolink 254 nm crosslinker (Thermo Fisher Scientific, Waltham, MA, USA) with an energy of 120 000 µJ.

Probe radiolabeling and filter hybridization were performed as described in (1). Hybridized filters were imaged with an Amersham™ Typhoon™Biomolecular Imager (GE Healthcare), and analyses were performed using HDFR software (Incogen, Williamsburg, NY, USA). Positive BAC clones detected by hybridization were validated individually by quantitative PCR (qPCR) amplification using the primer pairs used for probe synthesis and TME204 genomic DNA as control. qPCR amplified products were visualized using agarose gel electrophoresis and sequenced to confirm the specific amplification of targeted genes. BAC-end sequencing was used to compare and discriminate identified BAC clones. The insert size of the BAC clones was assessed using the FastNot I restriction enzyme and analyzed by pulsed field gel electrophoresis.

**BAC clone sequencing**

Individual BAC clone DNA were extracted using Nucleobond Xtra midi kit (Macherey-Nagel, Düren, Nordrhein-Westfalen). 2µg of each sample were used for the construction of a multiplexed SMRTbell® library by the INRAE-CNRGV. We followed the PacBio recommendations for Multiplexed Microbial Libraries preparation (PN 101-696-100) with some adjustments by using the SMRTbell Express Prep kit v2.0 (Pacific Biosciences, Menlo Park, CA, USA). The first enzymatic steps consist of removing single-stranded overhangs, repairing any DNA damage and polishing ends of the double stranded fragments and tailing with an A-overhang. Ligation with specific barcoded hairpin T-overhang adapters to both ends of the targeted double-stranded DNA (dsDNA) molecule was performed to create a closed, single-stranded circular DNA. A nuclease treatment was performed on each individual sample by using SMRTbell Enzyme Clean-up kit (Pacific Biosciences, Menlo Park, CA, USA). A size-selection with Blue-Pippin system (Sage Science, Beverly, MA, USA) to remove fragments less than 15Kb was done on pooled sample previously purified with 0.45X AMPure PB beads (Pacific Biosciences, Menlo Park, CA, USA). The size and concentration of the final library were assessed using the FemtoPulse system and the Qubit Fluorometer and Qubit dsDNA HS reagents Assay kit (Thermo Fisher Scientific, Waltham, MA, USA), respectively.

Sequencing primer v2 and Sequel DNA Polymerase 2.0 were annealed and bound, respectively to the SMRTbell library. The library was loaded on one SMRTcell at an on-plate concentration of 90pM using a diffusion loading. Sequencing was performed on the Sequel II system with a run movie time of 30 hours with 120 min pre-extension step and Software v9.0 (PacBio) by Gentyane Genomic Platform (INRAE-Clermont-Ferrand, France).

**BAC clone assembly**

The PacBio raw reads were corrected using SMRTLink_v9.0.0 with 8 passes, then demultiplexed. Residual *E. coli* reads were identified using BLAST+ 2.10.0 and removed using Seqfilter. The HiFi reads were filtered by identification of the vector sequences using cross_match and removed by custom Perl scripts. HiFi reads smaller than 15kb were filtered using Seqfilter, and then subsampled with SeqKit to obtain an estimated average assembly depth of 50x.  Assembly of the reads was performed with hifiasm-0.12. To validate the result, the length of the obtained contig were checked and BAC ends sequences were mapped with the extremities on the assembly using BLAST+ 2.10.0. HiFi Reads were remapped to the assembly and depth was obtained with samtools-1.8.

**Supplementary Figures**

**Supplementary figure 1. K-mer analysis reveal that genomes of cassava cultivars are diploid and highly heterozygous.**

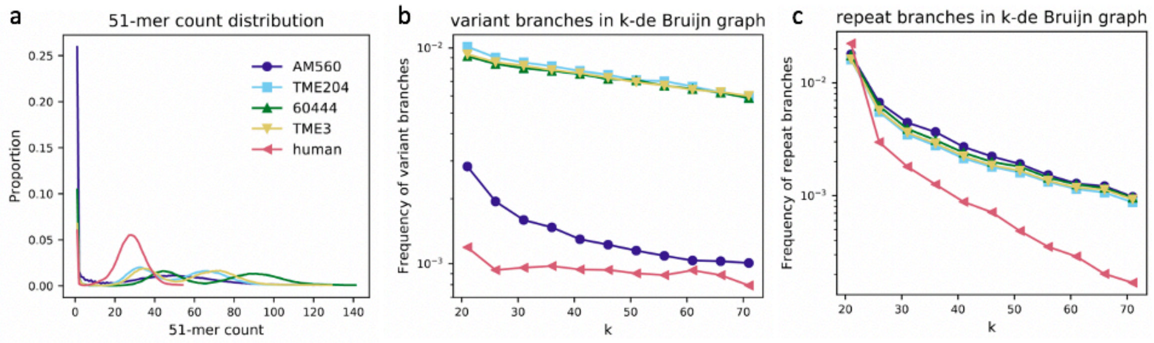


(a) Histogram of k-mer (length=51) coverage, which is the number of times a k-mer is observed in the Illumina reads (x-axis). The relative abundance of k-mers with a given coverage is plotted on the y-axis. For the African cassava TME204, TME3 and 60444 genomes, the k-mer coverage histograms are bi-modal, and the heterozygous peaks (at 35x for TME204/TME3, 45x for 60444) is as high as the homozygous peaks (at 70x for TME204/TME3 and 90x for 60444, respectively), indicating highly heterozygous genomes. For the inbred South-American cassava AM560 and human reference genomes, the histogram is single modal and dominated by the homozygous sequence peaks, indicating homozygous or nearly homozygous genomes. (b) Genome heterozygosity measured as the rate of variant branches (y-axis) in a de Bruijn graph as a function of k-mer length (x-axis). Approximately 1 in 100 vertices in the de Bruijn graphs of TME204, TME3 and 60444 has a variant-induced branch, corresponding to 1 SNP per 100 bp. The level of heterozygosity is 10 times higher than that in the human reference genome. The cassava reference genome was reconstructed from the inbred cassava cultivar AM560, in which the level of heterozygosity is similar to the level in the human reference genome. (c) Genome repetitiveness measured as the rate of repeat branches (y-axis) in a de Bruijn graph as a function of k-mer length (x-axis). The repetitiveness of all four cassava genomes is higher than that of the human genome when measured using k-mers ranging from 25 to 71 bp.

**Supplementary figure 2. Structural and phasing accuracy of the TME204 phased chromosome XII.**

a

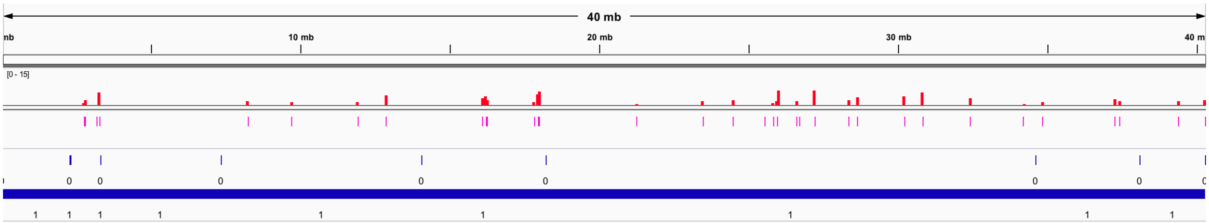


b

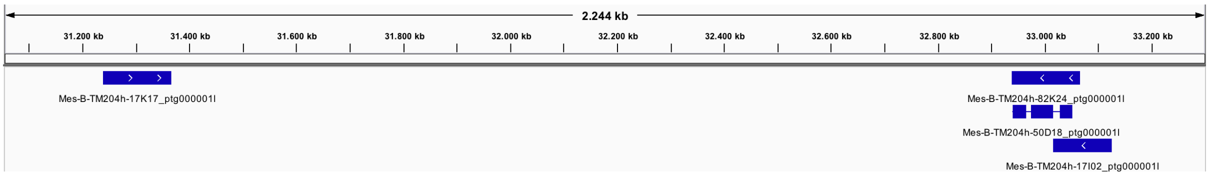


c

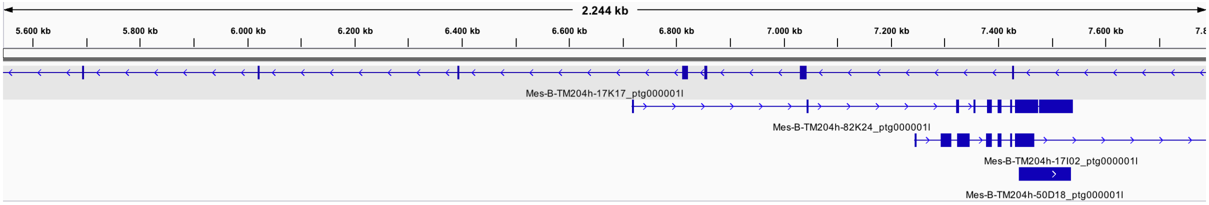

Each phased chromosome was assembled as one haplotig without sequencing gaps. (a) Structural accuracy of the phase 1 chromosome XII (minus-strand, h1tg00017l) measured by k-mers and CLRs. Potentially false duplications are highlighted by locations of k-mers present more than twice in the contig (as magenta bars) and their copy numbers (as red bars). Reliable blocks of assembled sequences with sufficient (≥10) PacBio CLR read support are shown as blue thick lines, labelled with 1 below. Potentially misassembled regions, identified by the lack of PacBio CLR read coverage (< 10), are highlighted as blue bars, labelled with 0 underneath. (b) Phasing accuracy of phase 1 chromosome XII (minus-strand, h1tg00017l) measured using BACs. (c) Phasing accuracy of phase 2 chromosome XII (plus-strand, h2tg00015l) measured using BACs. Three out of four sequenced BACs align continuously in full length with phase 1 chromosome XII. The fourth BAC (50D18) that cannot be aligned continuously with phase 1 aligns in full-length continuously with phase 2 chromosome XII.

**Supplementary figure 3. Distribution of uniquely and perfectly aligned genetic markers among TME204 haplotype 1 (a) and haplotype 2 (b) haplotigs**.

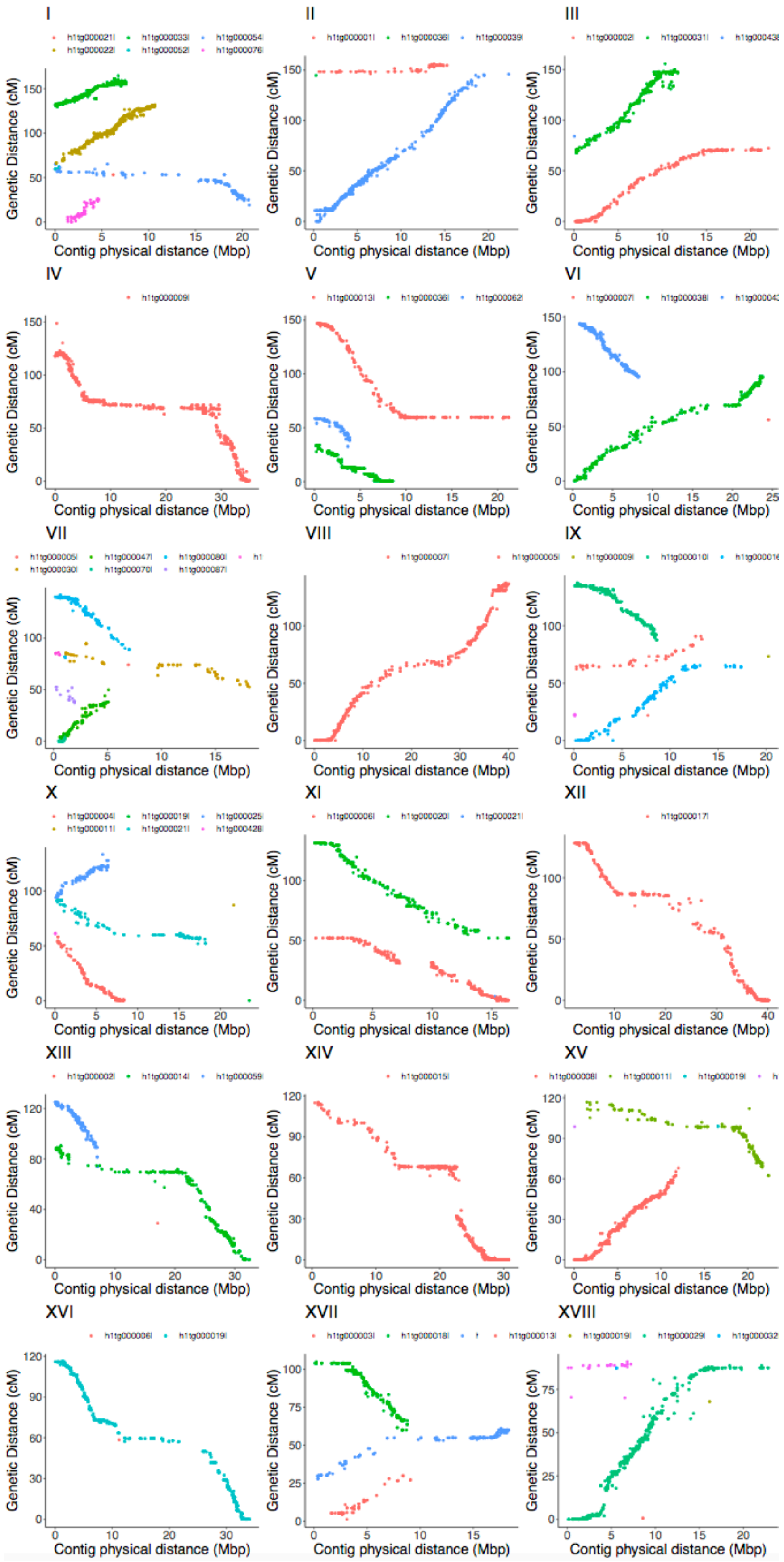

I
II
III
IV
V
VI
VII
VIII
IX
X
XI
XII
XIII
XIV
XV
XVI
XVII
XVIII
Genetic Distance (cM)
Contig physical distance (Mbp)

b

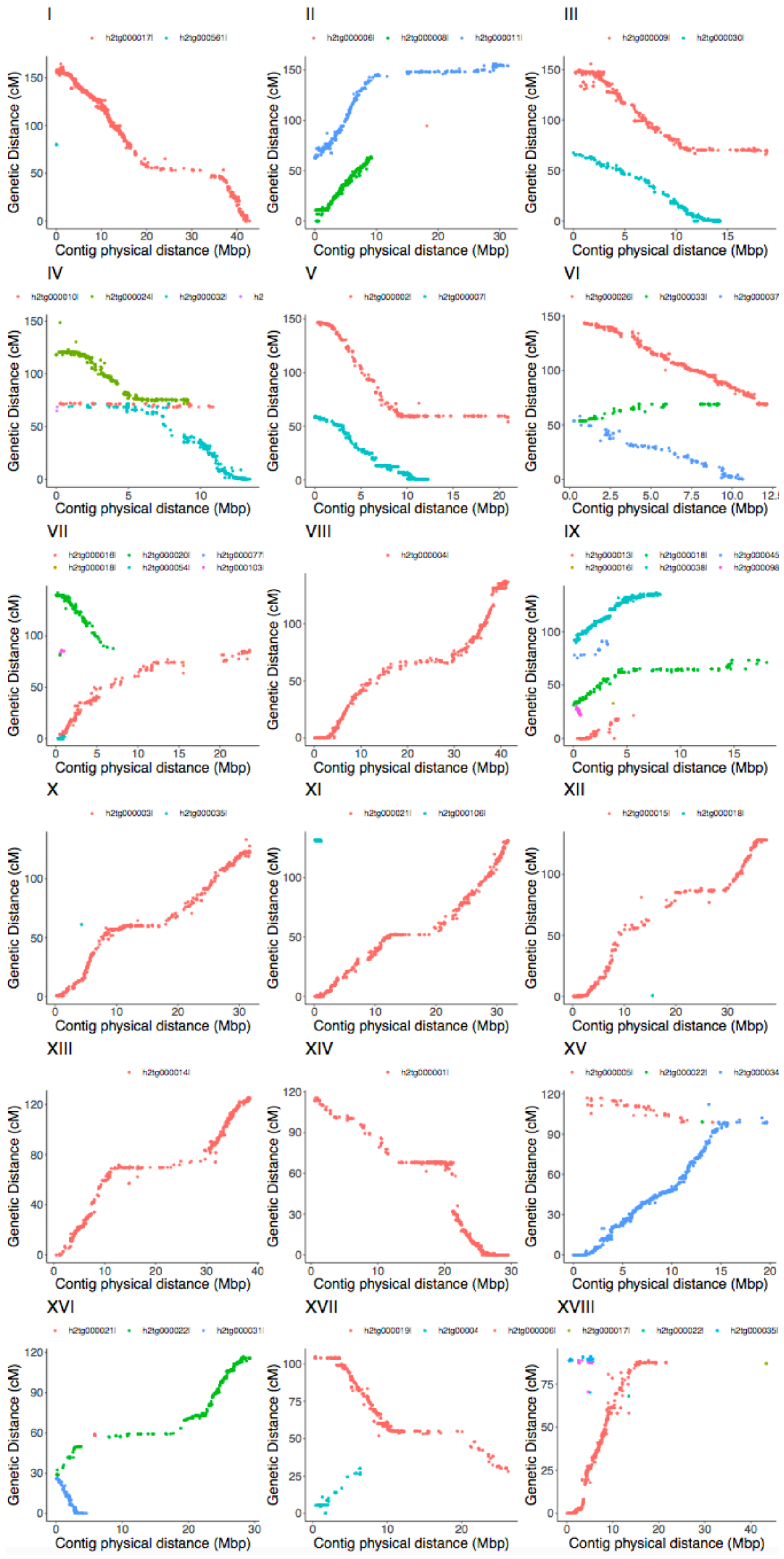
I
h2tg000017l
h2tg000561l
II
h2tg000006l
h2tg000008l
h2tg000011l
III
h2tg000009l
h2tg000030l
IV
h2tg000010l
h2tg000024l
h2tg000032l
V
h2tg000002l
h2tg000007l
VI
h2tg000026l
h2tg000033l
VII
h2tg000016l
h2tg000020l
h2tg000077l
h2tg000018l
h2tg000054l
h2tg000103l
VIII
h2tg000004l
IX
h2tg000013l
h2tg000018l
h2tg000016l
h2tg000038l
X
h2tg000003l
h2tg000035l
XI
h2tg000021l
h2tg000106l
XII
h2tg000015l
h2tg000018l
XIII
h2tg000014l
XIV
h2tg000001l
XV
h2tg000005l
h2tg000022l
XVI
h2tg000021l
h2tg000022l
h2tg000031l
XVII
h2tg000019l
h2tg000006l
XVIII
h2tg000017l
h2tg000022l
h2tg000035l
Genetic Distance (cM)
Contig physical distance (Mbp)

The presence of phased chromosomes and chromosome pairs can be visually identified at the contig level without sequencing gaps. Most of the 18 chromosome pairs consist of only a few haplotigs. Genetic distance (cM) on the y-axis is derived from the genetic map. Physical distance (Mbp) on the x-axis is derived from the uniquely aligned positions in the TME204 haplotigs. Each dot is a genetic marker. Different colors represent different haplotigs.

**Supplementary figure 4. Haplotigs of the mitochondrial genome and chromosomal distribution of numt's in the TME204 diploid genome.**

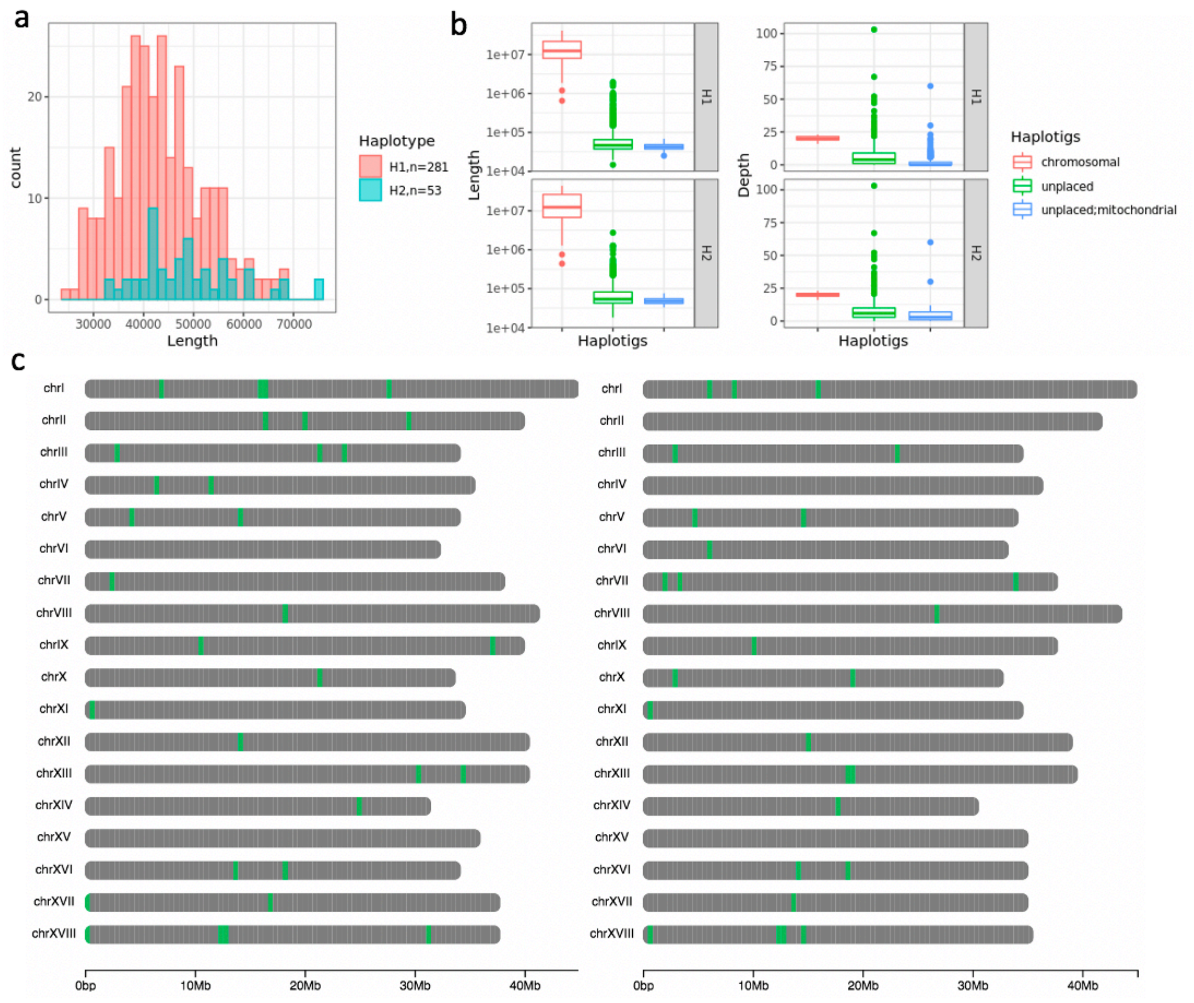


(a) Total counts and length distributions of mitochondrial haplotigs in TME204 H1 and H2 assemblies. (b) Length and depth of coverage for chromosome anchored haplotigs, unanchored mitochondrial haplotigs, and the other unanchored haplotigs. (c) Chromosomal distribution of numt's in TME204 H1 and H2 assemblies.

**Supplementary figure 5. Repeat landscape of cassava TME204 diploid genome.**

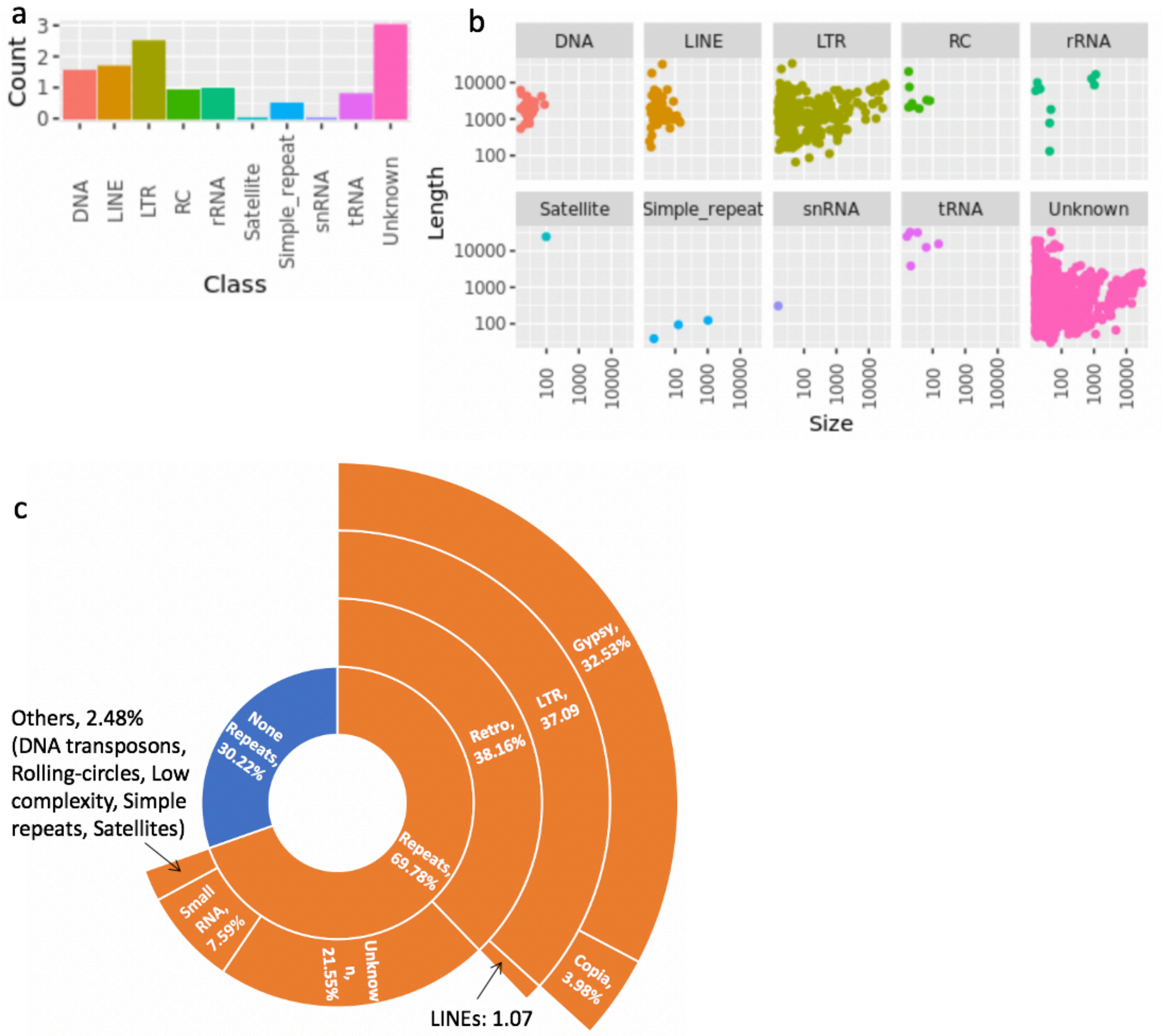

(a) Number of *de novo* predicted repeat families per repeat class. (b) Length and copy number of *de novo* predicted repeat families, grouped by repeat classes. (c) Repeat landscape of cassava TME204 H1 assembly.

**Supplementary figure 6. Haplotype specific formation of incomplete transcripts in the TME204 genome.**

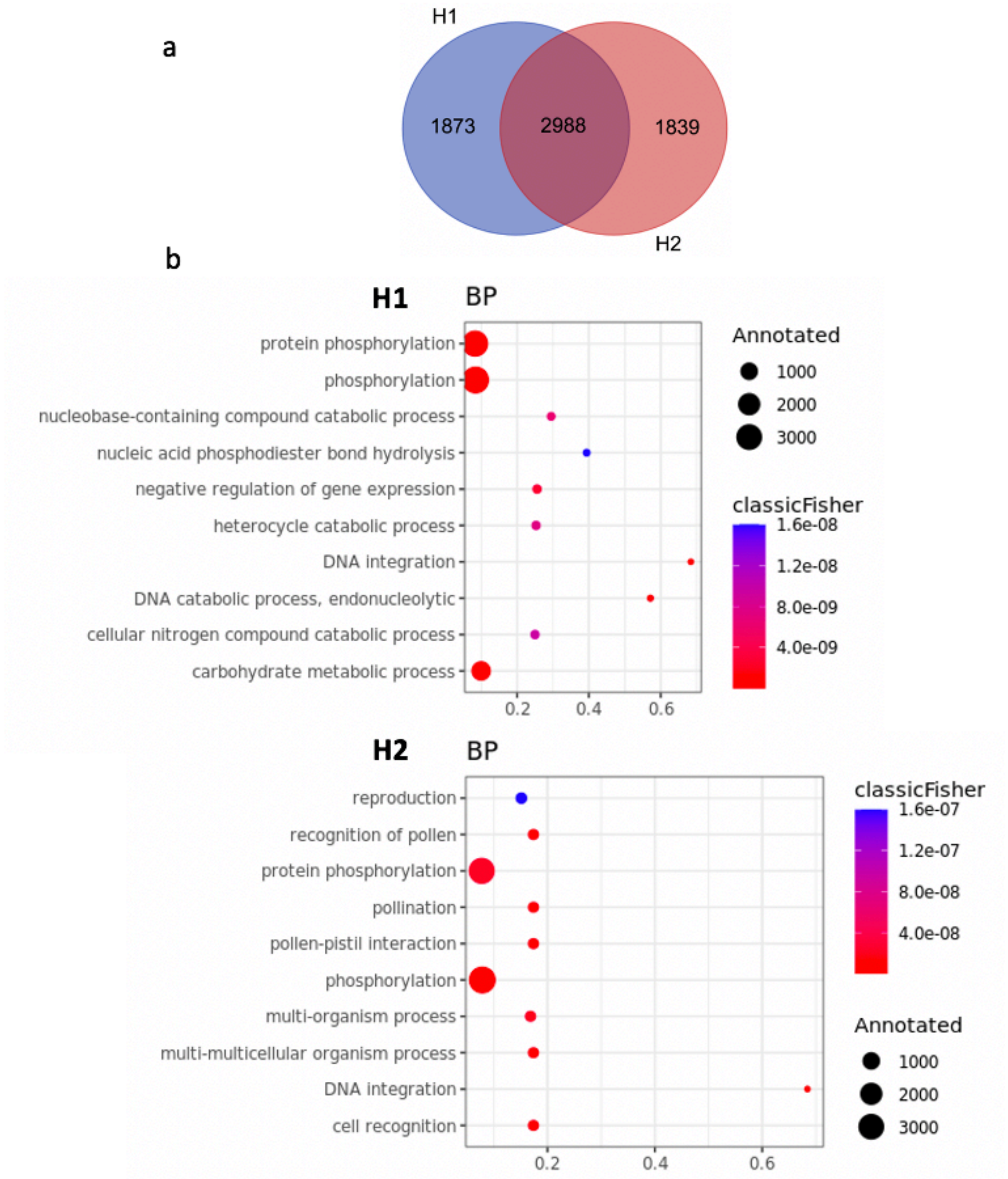


(a) Venn diagram showing the number of haplotype specific incomplete transcripts and those shared by both haplotypes. (b) Enriched biological process (BP) GO terms identified in sets of H1-specific and H2-specific disrupted transcripts.

**Supplementary figure 7. Structural category of PacBio Iso-Seq transcripts aligned to the disrupted genes in the TME204 H2 assembly.**

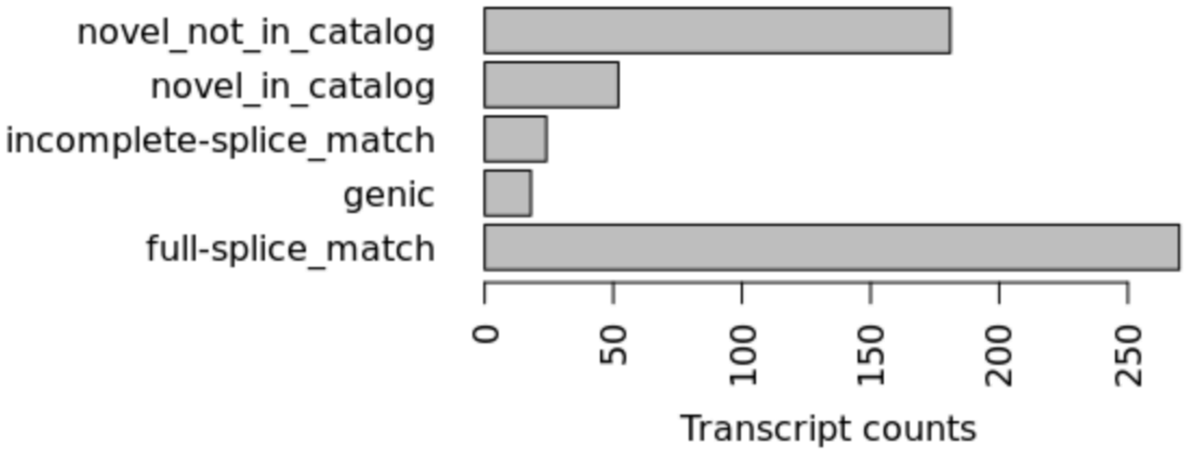


**Supplementary figure 8. Functional enrichment analysis of TME204 novel genes identified with Iso-Seq transcripts.**

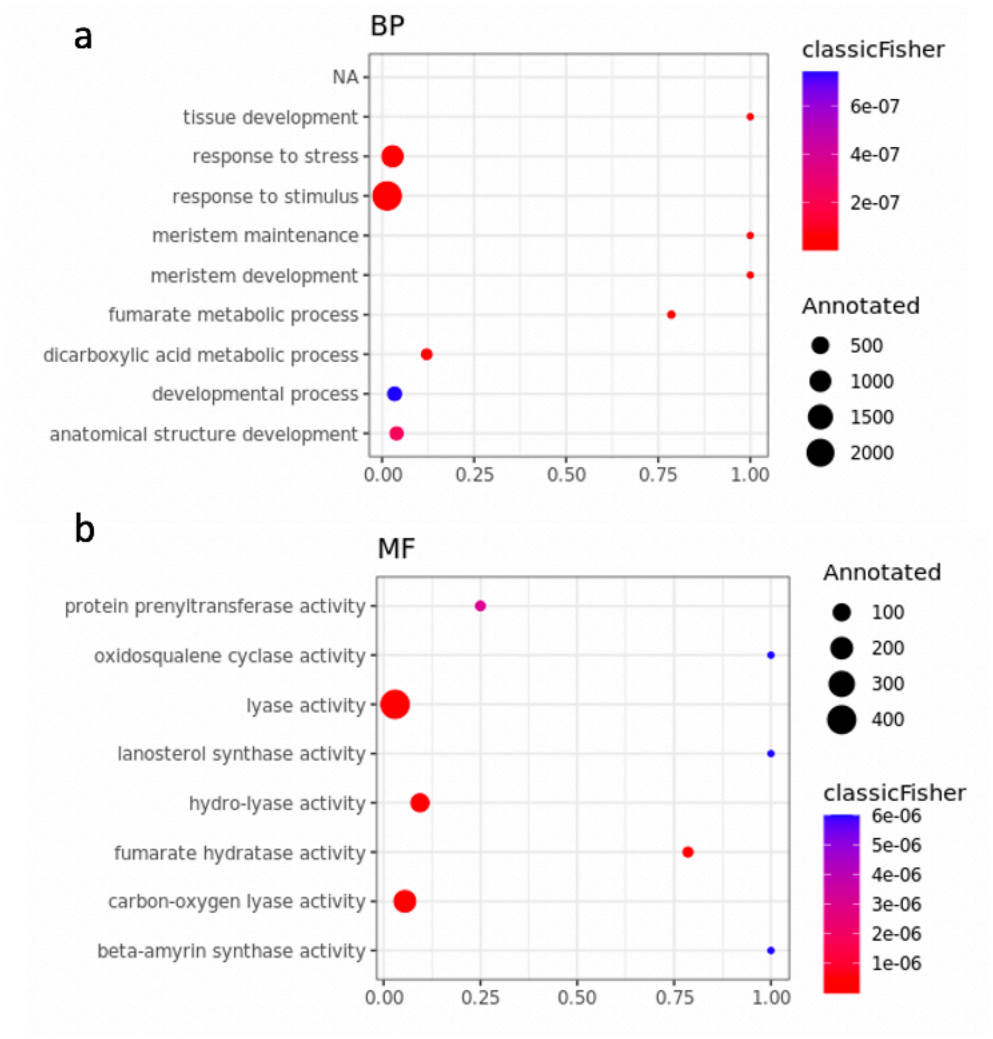


(a) Enriched GO terms in the category of biological process (BP). (b) Enriched GO terms in the category of molecular function (MF).

**Supplementary figure 9. Classifications and size distributions of small and large indels identified by Assemblytics analysis of reliable contig alignments between cassava genomes.**

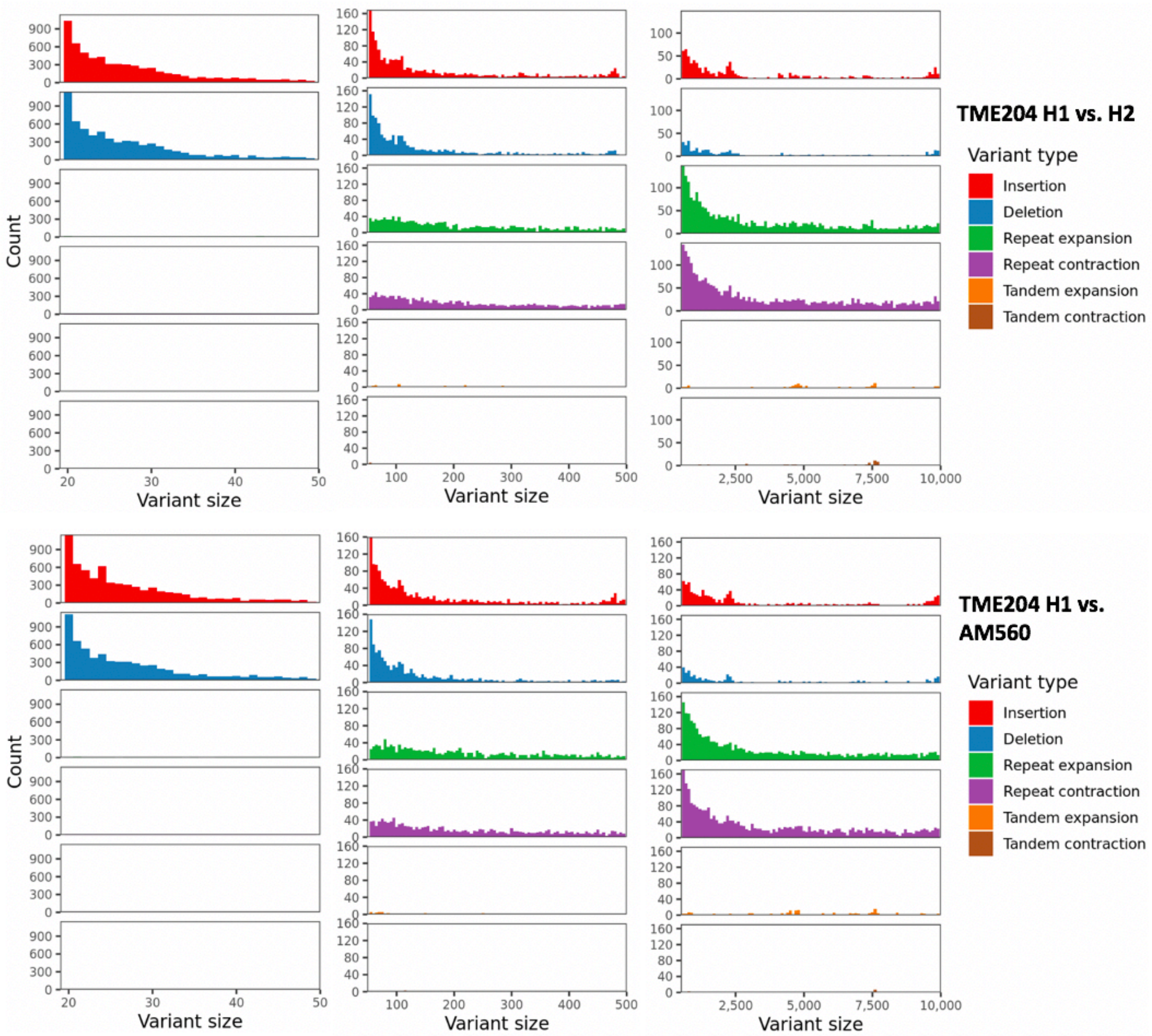


(a) Indels between TME204 H1 and H2 haplotigs. (b) Indels between TME204 H1 haplotigs and AM560 contigs.

**Supplementary tables**

**Supplementary table 1. Assembly statistics for different cassava TME204 primary and alternate genome assemblies based on PacBio CLR and HiFi reads.**

| Contig set | Read type | Assembler | Size (Mbp) | N50 (Mbp) | BUSCO complete (%) | BUSCO duplicate (%) |
|---|---|---|---|---|---|---|
| Primary | CLR | Falcon | 1176.669 | 1.261 | 95.3 | 46.5 |
| | HiFi | Falcon | 995.112 | 3.841 | 91.2 | 14.7 |
| | | Hifiasm | 922.872 | 29.016 | 96.8 | 15.6 |
| | | IPA | 1014.359 | 4.698 | 96.8 | 43.5 |
| Alternate | CLR | Falcon | 58.250 | 0.267 | 18.1 | 0.7 |
| | HiFi | Falcon | 185.785 | 0.438 | 38.6 | 1.9 |
| | | Hifiasm | 617.610 | 0.274 | 83.8 | 14.7 |
| | | IPA | 223.050 | 0.647 | 51.6 | 2.1 |

**Supplementary table 2. Assembly statistics for different cassava TME204 combined assemblies of mixed-haplotypes based on PacBio CLR and HiFi reads.**

| Read type | CLR | HiFi | HiFi | HiFi | HiFi |
|---|---|---|---|---|---|
| Assembler | Falcon | Falcon | HiCanu | Hifiasm | IPA |
| Version | pb-assembly 0.06 | pb-assembly 0.08 | 2.0 | 0.7 | 1.05 |

| | | | | | |
|---|---|---|---|---|---|
| Assembled sequences | Primary and associated contigs | Primary and associated contigs | Contigs of all resolved Alleles | Primary and purged contigs, phased haplotigs | Primary and phased haplotigs |
| Number of bases (Gbp) | 1.235 | 1.181 | 1.484 | 1.540 | 1.237 |
| Contig N50 (Mbp) | 1.189 | 2.803 | 8.131 | 6.464 | 3.363 |
| Contig NG50 (Mbp) | 2.775 | 5.288 | 23.781 | 33.087 | 6.302 |
| Largest contig (Mbp) | 7.773 | 18.554 | 42.800 | 42.956 | 17.899 |
| BUSCO complete (%) | 95.6 | 92.5 | 96.8 | 96.3 | 96.5 |
| BUSCO duplicate (%) | 54.0 | 28.1 | 82.1 | 82.1 | 76.5 |

**Supplementary table 3. Cassava TME204 genome assembly benchmarking results for PacBio CLR and HiFi reads, measured by alignments and k-mer analysis of Illumina PE reads.**

| Read types | CLR | HiFi | | | |
|---|---|---|---|---|---|
| Assemblers | Falcon | Falcon | HiCanu | Hifiasm | IPA |
| Assembled sequences | All resolved alleles combined | | | | |

| | | | | | |
|---|---|---|---|---|---|
| Mapping rate (%) | 98.29 | 98.21 | 99.88 | 99.88 | 98.21 |
| Properly paired (%) | 94.75 | 95.41 | 99.33 | 99.32 | 95.12 |
| Singletons (%) | 0.34 | 0.41 | 0.02 | 0.02 | 0.42 |
| Mate mapped to a different contig (%) | 3.10 | 2.30 | 0.48 | 0.48 | 2.61 |
| Other wrongly pairs (%) | 0.1 | 0.09 | 0.05 | 0.06 | 0.06 |
| General error rate (%) | 0.68 | 0.59 | 0.21 | 0.21 | 0.38 |
| Total k-mer in the assembly | 1,234,874,650 | 1,180,852,607 | 1,483,627,597 | 1,540,334,405 | 1,237,344,741 |
| K-mer not in Illumina data | 38,973,344 | 39,754,338 | 2,157,169 | 652,749 | 1,690,846 |
| K-mer survival rate (%) | 99.840 | 99.829 | 99.993 | 99.997 | 99.993 |
| Consensus quality (QV) | 27.95 | 27.67 | 41.38 | 46.74 | 41.65 |
| K-mer completeness | 93.20 | 94.96 | 98.40 | 98.40 | 97.54 |

**Supplementary table 4. Cassava TME204 BAC-to-haplotig alignments.**

| BAC | Length (bp) | Haplotype 1 | Haplotype 2 |
|---|---|---|---|

| | | Haplotig | Number aligned blocks | Number matches (bp) | Haplotig | Number aligned blocks | Number matches (bp) |
|---|---|---|---|---|---|---|---|
| 17I02 | 110289 | h1tg000017l | 2[a] | 110288 | h2tg000015l | 433 | 76783 |
| 17K17 | 128326 | h1tg000017l | 1 | 128326 | h2tg000015l | 288 | 44218 |
| 50D18 | 95702 | h1tg000017l | 253 | 74458 | h2tg000015l | 2[a] | 95701 |
| 82K24 | 126840 | h1tg000017l | 1 | 126835[b] | h2tg000015l | 414 | 88170 |

[a] The two-block alignment was due to one insertion (1bp) in the assembled BAC sequence.

[b] A 5 bp sequence at the 3' end of the assembled BAC sequence was not aligned. Manual inspection revealed that they belong to the cloning vector.

**Supplementary table 5. Hi-C scaffolds in cassava TME204 haplotype 2 assembly.**

| Scaffold ID | Scaffold length (Mbp) | Haplotig ID | Chromosome placement by the genetic map |
|---|---|---|---|
| 1 | 113 | h2tg000005l | XV |
| | | h2tg000034l | XV |
| | | h2tg000017l | I |
| | | h2tg000006l | XVIII |
| | | h2tg000046l | XVIII |
| | | h2tg000035l | XVIII |
| 3 | 41 | h2tg000008l | II |
| | | h2tg000011l | II |
| 6 | 35 | h2tg000032l | IV |
| | | h2tg000010l | IV |

| | | | |
|---|---|---|---|
| | | h2tg000024l | IV |
| 7 | 34 | h2tg000022l | XVI |
| | | h2tg000031l | XVI |
| 8 | 34 | h2tg000019l | XVII |
| | | h2tg000047l | XVII |
| 9 | 34 | h2tg000009l | III |
| | | h2tg000030l | III |
| 10 | 33 | h2tg000007l | V |
| | | h2tg000002l | V |
| 12 | 32 | h2tg000037l | VI |
| | | h2tg000033l | VI |
| | | h2tg000026l | VI |
| 37 | 0.4 | h2tg000089c | NA |
| | | h2tg000642l | NA |
| 102 | 0.1 | h2tg000530l | NA |
| | | h2tg000628l | NA |
| 159 | 0.1 | h2tg000514l | NA |
| | | h2tg000694l | NA |
| 168 | 0.1 | h2tg000495l | NA |
| | | h2tg000607l | NA |

**Supplementary table 6. Assembled sequences with nuclear mitochondrial pseudogene regions (numt's) in the TME204 genome.**

| Sequences | H1 | H2 |
|---|---|---|

| | | |
|---|---|---|
| Chromosome with numt’s | 16 | 13 |
| Unanchored haplotigs with numt’s | 592 | 265 |

**Supplementary table 7. Gene synteny and inversions identified using orthologous gene pairs including disrupted genes and genes with more degenerated sequences in TME204 assemblies.**

| | TME204 H1 vs. TME204 H2 | AM560 vs. TME204 H2 |
|---|---|---|
| Percentage (%) of collinear genes | 97.15 | 97.10 |
| Inversions | 10 | 12 |

**Supplementary table 8. TME204 PacBio Iso-Seq reads and high quality (HQ [a]) transcripts.**

| Tissue | PacBio barcode name | Polymerase reads | Accession number | HQ transcripts |
|---|---|---|---|---|
| Leaf | bc1008_5p--bc1008_5p | 180,025 | ERR5489420 | 23,103 |
| Stem | bc1012_5p--bc1012_5p | 290,953 | ERR5489421 | 38,333 |
| Root | bc1018_5p--bc1018_5p | 301,754 | ERR5489422 | 36,147 |

[a] Accuracy 99.9% and above

**Supplementary table 9. Validation and improvement of TME204 gene and transcript annotation using PacBio Iso-Seq HQ transcripts.**

| Validated Genes | Validated Transcripts | H1 | H2 |
|---|---|---|---|
| Lift over genes | | 15,058 | 15,030 |
| | Lift over transcripts | 18,624 | 18,593 |
| | Novel transcripts (Fusion transcripts) | 3,881 (240) | 3,841 (251) |
| | Fusion genes | 268 | 275 |
| | Genomic loci of fusion genes | 132 | 135 |
| Novel genes | | 426 | 408 |
| | Novel transcripts | 4,335 | 4,273 |

**Supplementary table 10. Reanalysis of a published TME204 Illumina RNA-seq dataset (2).**

| Accession numbers | Tissue type |
|---|---|
| SRR3629818, SRR3629835, SRR3629853 | Leaf |
| SRR3629824, SRR3629842, SRR3629859 | Stem |
| SRR3629843 | Storage root |
| SRR3629845 | Fibrous root |
| SRR3629837 | Midvein |
| SRR3629838 | Petiole |
| SRR3629840 | Lateral bud |
| SRR3629850 | SAM |
| SRR3629851 | RAM |

**Supplementary table 11. Primer sequences used to amplify the probes used for BAC screening.**

| BAC | Primer Name | Primer Sequence |
|---|---|---|

| | | |
|---|---|---|
| 17K17 | CMD396 | TTGCAGGAGGACACACCATTGG |
| | CMD397 | ACCTTGGCAGCCAGTGAGG |
| 17I02 | CMD410 | GACCGCTTTCACTGGTACCCAC |
| | CMD411 | GCAGTAGAACTCCCACGTCCGT |
| 50D18, 82K24 | CMD408 | ATGCTAGACCGCTGTGCATGC |
| | CMD409 | GTGGAGAACTGGCCTGCTCAG |
